# Chemical Proteomics of Residual Acute Myeloid Leukemia Cells Reveals Therapeutic Vulnerabilities

**DOI:** 10.64898/2026.09.22.752285

**Authors:** Elham Gholizadeh, Uladzislau Vadadokhau, Mücahit Varli, Danilo Ritz, Komal Kumar Javarappa, Mika Kontro, Blassan George, Raphael Itzykson, Markku Varjosalo, Risto Renkonen, Esko Kankuri, Paul A. Haynes, Amir A. Saei, Mohieddin Jafari

**Affiliations:** Department of Pharmacology, Faculty of Medicine, University of Helsinki, Helsinki, Finland; Institute for Molecular Medicine Finland (FIMM), HiLIFE, University of Helsinki, Helsinki, Finland; Department of Microbiology, Tumor and Cell Biology, Karolinska Institutet, Stockholm, Sweden; Biozentrum, University of Basel, Basel, Switzerland; Department of University Sophisticated Instrumentation Centre (USIC), JSS AHER, Mysuru, Karnataka, India; Laser Research Centre, Faculty of Health Sciences, University of Johannesburg, P.O. Box 17011, Doornfontein 2028, South Africa; Institut de Recherche Saint-Louis, UMR1342, Université Paris Cité, INSERM, Paris, France; Institute of Biotechnology and Helsinki Institute of Life Science, University of Helsinki, 00014 Helsinki, Finland; Faculty of Medicine, University of Helsinki and Helsinki University Hospital, Helsinki, Finland; School of Natural Sciences, Macquarie University, North Ryde, New South Wales 2109, Australia; Faculty of Medicine and Health Technology, Tampere University and TAYS Cancer Center, Tampere, Finland; Tampere Institute for Advanced Study, Tampere University, Tampere, Finland

## Abstract

Venetoclax combined with azacitidine has improved treatment outcomes in acute myeloid leukemia (AML), yet relapse and treatment persistence remain major clinical challenges. To investigate proteomic mechanisms associated with venetoclax–azacitidine response and adaptation, we applied a multi-layer combinatorial proteome integral solubility/stability alteration analysis (CoPISA) strategy in SKM-1 AML cells. Cells were treated with venetoclax, azacitidine, their combination, or vehicle control and profiled across four orthogonal layers: short-term lysate CoPISA, short-term intact-cell CoPISA, long-term intact-cell CoPISA after 5 days of treatment, and long-term expression proteomics of surviving cells. The venetoclax–azacitidine combination induced treatment-specific protein solubility and abundance changes that were not fully reproduced by either single agent. Long-term surviving cells displayed extensive proteomic remodeling, consistent with the emergence of an adaptive drug-tolerant state, although contributions from pre-existing resilient cell populations cannot be excluded. Integration of short- and long-term solubility changes with abundance remodeling revealed distinct adaptive regimes, including retained biochemical targets, dosage-compensated targets, sensitive-state-specific targets, and remodeled adaptive targets. This framework prioritized candidate resistance-associated proteins, including NRP2, RPL18, PLP2, RPS28, and NOTCH1. NRP2 emerged as a top candidate across all proteomics layers, consistently showing increased abundance. Overall, this study shows that combining CoPISA with expression proteomics can resolve temporally distinct proteomic states of venetoclax–azacitidine response and identify candidate adaptive vulnerabilities in AML.

## Introduction

Acute myeloid leukemia (AML) remains a clinically challenging hematological malignancy because durable remission is frequently limited by treatment persistence and relapse. The combination of the BCL2 inhibitor venetoclax with the hypomethylating agent azacitidine has improved outcomes for older or intensive-chemotherapy-ineligible patients, establishing venetoclax–azacitidine (VA) as a major therapeutic option in AML [1]. Nevertheless, responses are rarely curative, and relapse after venetoclax-based therapy is associated with poor outcomes and limited effective treatments [2–5]. Defining how AML cells survive VA pressure is therefore essential for identifying new therapeutic vulnerabilities.

Historically, AML relapse has been explained by clonal evolution, selection of pre-existing resistant subclones, and persistence of leukemic stem cell populations. Some evidence, however, indicates that treatment failure can also arise through non-genetic cellular adaptation. Drug-tolerant persister (DTP) cells represent a transient state in which a minority of cancer cells survive acute drug exposure without initially acquiring stable resistance mutations [6]. These cells often display slow proliferation, altered apoptotic priming, metabolic rewiring, stress-response activation, epigenetic remodeling, and context-dependent changes in cell identity. After removal of drug pressure, DTP cells can regain drug sensitivity, distinguishing them from permanently resistant cells; however, under continued treatment, they may provide a reservoir from which stable acquired resistance can emerge [6,7].

In AML, several studies now support the existence of treatment-induced persister-like states. Chemotherapy-resistant residual AML cells have been shown to rely on oxidative metabolism rather than being uniformly enriched for classical leukemic stem cell markers [8]. More recently, AML DTP cells surviving daunorubicin and cytarabine (Ara-C) were shown to transiently increase plasma membrane rigidity, reducing chemotherapy uptake and promoting short-term survival while simultaneously increasing susceptibility to T-cell-mediated killing [9]. Together, these findings indicate that AML persistence is not a single fixed phenotype but a dynamic and potentially targetable adaptive state.

Venetoclax-based treatment provides a clinically relevant context in which such adaptive states may emerge. Venetoclax induces apoptosis primarily in BCL2-dependent AML cells by promoting mitochondrial outer membrane permeabilization, cytochrome c release, and subsequent caspase activation. Sustained exposure to venetoclax can select for cells with reduced dependence on BCL2 and increased reliance on alternative anti-apoptotic proteins, including MCL1 and BCL-xL, as well as cells with altered mitochondrial metabolism and apoptotic regulation [10]. These mechanisms have been primarily described in the context of venetoclax resistance; however, the transition from drug-induced persistence to stable resistance may involve intermediate cellular states that retain substantial biochemical consequences of drug exposure. Importantly, venetoclax-resistant AML cells can retain sublethal apoptotic and cellular stress responses following venetoclax exposure, indicating that loss of drug sensitivity does not necessarily imply complete loss of drug-induced cellular perturbation [10]. This distinction between drug sensitivity, transient persistence, and established resistance highlights the need to characterize protein-level changes that occur both early during drug exposure and after prolonged treatment.

A central obstacle in studying DTP biology is that these cells are defined primarily by behavior rather than by a stable genetic lesion. No universal DTP biomarker exists, and DTP-associated features vary according to cancer type, drug exposure, treatment duration, and experimental model [6,7]. Methods that capture both early drug-induced protein engagement and later adaptive proteome remodeling are therefore needed. Solubility-based thermal proteome profiling approaches, including the proteome integral solubility alteration (PISA) assay, can detect treatment-induced changes in protein stability and solubility at the proteome scale without requiring chemical modification of the drug [11–13]. The recently developed combinatorial proteome integral solubility/stability alteration analysis (CoPISA) extends this strategy to drug combinations, enabling systematic identification of protein solubility responses induced by single agents and combination treatments [14]. CoPISA can be performed in either cell extracts or intact living cells, with each format capturing different aspects of drug-induced protein-state changes. In cell extracts, drug exposure occurs after cell disruption, minimizing the influence of membrane transport, intracellular metabolism, compartmentalization, and cellular feedback mechanisms. This simplified biochemical setting may therefore favor the detection of direct or proximal drug-induced protein-state changes. In contrast, intact-cell CoPISA preserves the cellular environment during drug exposure, enabling the detection of changes associated not only with target engagement but also with cellular responses such as protein-complex remodeling, subcellular relocalization, post-translational regulation, stress signaling, apoptotic priming, and early adaptive remodeling [15–18]. Thus, comparison of extract-based and intact-cell CoPISA was used to separate extract-detectable protein-state changes from responses that depend on intact-cell biology [19,20].

In this study, we applied a multi-layer proteomic strategy to investigate VA response and adaptation in SKM-1 AML cells (**Fig. 1**). To capture distinct temporal and cellular states of drug response, we integrated four orthogonal proteomic layers: short-term cell extract CoPISA (*SC*), short-term living-cell CoPISA (*SL*), long-term living-cell CoPISA (*LL*), and long-term expression proteomics (*LE*). We define significantly altered protein sets within each layer as 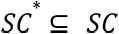 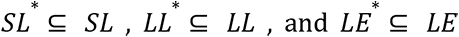, and further define proteins with increased abundance in resistant or DTP-enriched states as 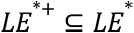, whereas proteins below the predefined abundance cutoff were defined as 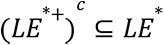. Within this framework, we speculate that protein behavior across these layers may reveal distinct aspects of drug response and resistance adaptation. Proteins detected in both short- and long-term CoPISA without increased abundance may be consistent with drug-engaged targets that remain biochemically accessible without evidence of compensatory expression (R_1_). Proteins that persist across CoPISA layers and are also increased in abundance in resistant cells may indicate functional nodes under sustained drug pressure, potentially reflecting compensatory mechanisms associated with resistance (R_2_). In contrast, proteins identified only in short-term CoPISA and absent in long-term measurements without corresponding expression changes may represent interactions characteristic of the sensitive state that are no longer maintained following the acquisition of resistance (R_3_). Finally, short-term targets that are not retained in long-term CoPISA yet become increased in abundance suggest a more complex adaptive mechanism involving both compensatory expression and potential target remodeling, such as post-translational modification (PTM) [16] or altered complex formation, leading to reduced effective drug engagement (R_4_). Formally, these four regimes can be defined as follows (**Fig. 1B**):

- Sensitive–Persistent common targets:

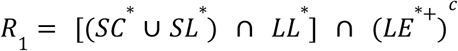
- Persistent-adaptive dosage compensation:

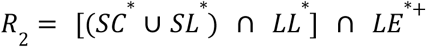
- Sensitive-specific targets:

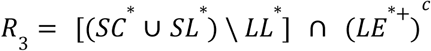
- Adaptive remodeled targets:

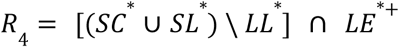

**Figure 1:**
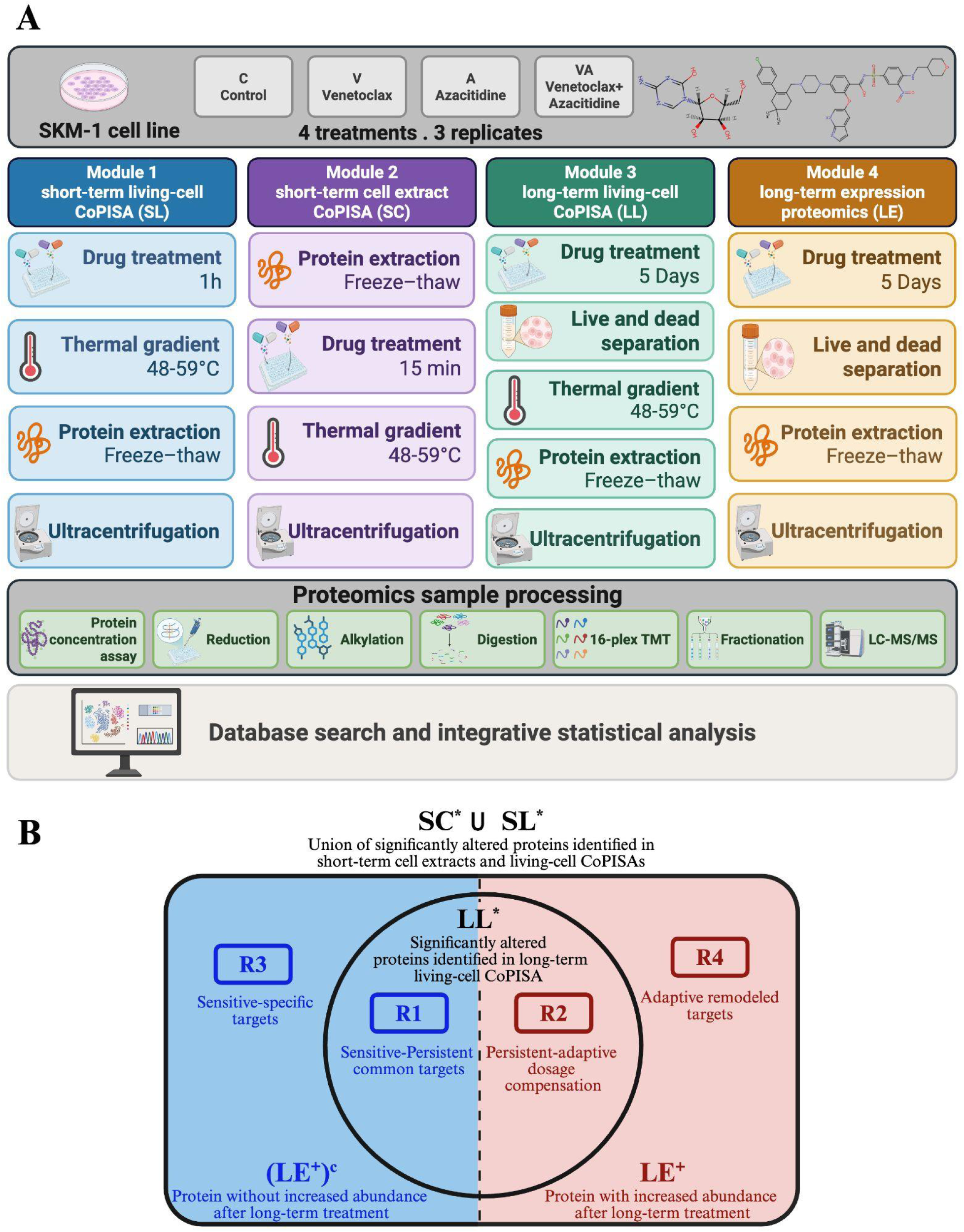
Integrated proteomic workflow and classification framework for identifying treatment-responsive targets in AML cells. **A)** SKM-1 AML cells were treated with DMSO (control), venetoclax, azacitidine, or combinations of venetoclax plus azacitidine and profiled across orthogonal proteomics modalities, including short-term living-cell CoPISA (SL), short-term cell extract CoPISA (SC), long-term living-cell CoPISA (LL), and long-term expression proteomics (LE). Soluble protein fractions were processed by mass spectrometry and integrated using differential solubility analysis, permutation-based robustness assessment, and cross-platform meta-analysis to prioritize treatment-responsive proteins. **B)** Schematic representation of the hierarchical classification of sensitive-responsive targets. The sensitive-responsive target universe is first partitioned into persistent 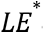 and non-persistent targets, and each subset is subsequently divided according to adaptive remodeling status 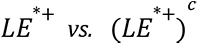. This defines four mutually exclusive target classes: **R1**, sensitive–persistent common targets; **R2**, persistent-adaptive dosage compensation targets; **R3**, sensitive-specific targets; and **R4**, adaptive remodeled targets.

Collectively, this framework allows us to disentangle early drug-target engagement from long-term resistance adaptation and to prioritize proteins that are not only initially drug-responsive but also actively remodeled in resistant states. We propose that *R*_2_ and *R*_4_ capture classes of compensatory resistance mediators, where cancer cells maintain survival through combined strategies of target modulation and expression-level adaptation, thereby highlighting actionable nodes for combination therapeutic intervention.

## Results

### CoPISA workflow and multi-layer proteomic response to VA treatment

To investigate how the venetoclax-azacitidine combination affects AML cells and to identify potential mechanisms of resistance, we applied our recently developed CoPISA assay workflow to the SKM-1 cell line. Cells were treated with venetoclax (V), azacitidine (A), the VA combination, and DMSO control (C), and the workflow was performed under both short-term and long-term treatment conditions (**Fig. 1**).

CoPISA measures drug-induced changes in protein solubility or stability by calculating the total soluble abundance of each protein across a thermal gradient, represented as the area under the melting curve (Sm). This allows detection of proteins whose solubility changes after drug treatment without requiring full melting-curve fitting. In this study, CoPISA was applied to both cell extracts and intact living cells, enabling comparison of direct drug-associated effects in lysates with broader cellular responses occurring in living cells.

As shown in **Fig. 1A**, the workflow included both short-term and long-term treatment modules. In the short-term module, CoPISA was performed in both cell extracts (SC) and living cells (SL) to capture early drug-induced solubility changes after treatment with venetoclax, azacitidine, the VA combination, or DMSO control. In the long-term module, cells were treated for 5 days, followed by two parallel analyses: long-term living-cell CoPISA (LL) and long-term expression proteomics (LE).

Across all TMT-based proteomics datasets, a total of 5,578 unique proteins and 26,447 unique peptides were identified after quality-control filtering. Reverse-sequence identifications and potential contaminants were excluded, and peptide-spectrum evidence with no detectable corrected reporter-ion intensity across all TMT channels was removed. Individual datasets showed broad proteome coverage, ranging from 3,176 to 4,735 proteins and from 9,366 to 17,057 peptides, supporting sufficient identification depth for downstream differential solubility and expression analyses (**Supplementary Fig. S1**).

### Multi-layer proteomic profiling reveals distinct short- and long-term drug responses

To determine the appropriate time point for long-term treatment, SKM-1 cell viability was monitored daily using the trypan blue exclusion assay after treatment with venetoclax, azacitidine, the VA combination, or DMSO control. As shown in **Fig. 2A**, control cells maintained high viability throughout the experiment, remaining above ∼80% across the 5-day period. Azacitidine alone caused only a moderate decrease in viability, while venetoclax and VA produced a stronger reduction.

**Figure 2.**
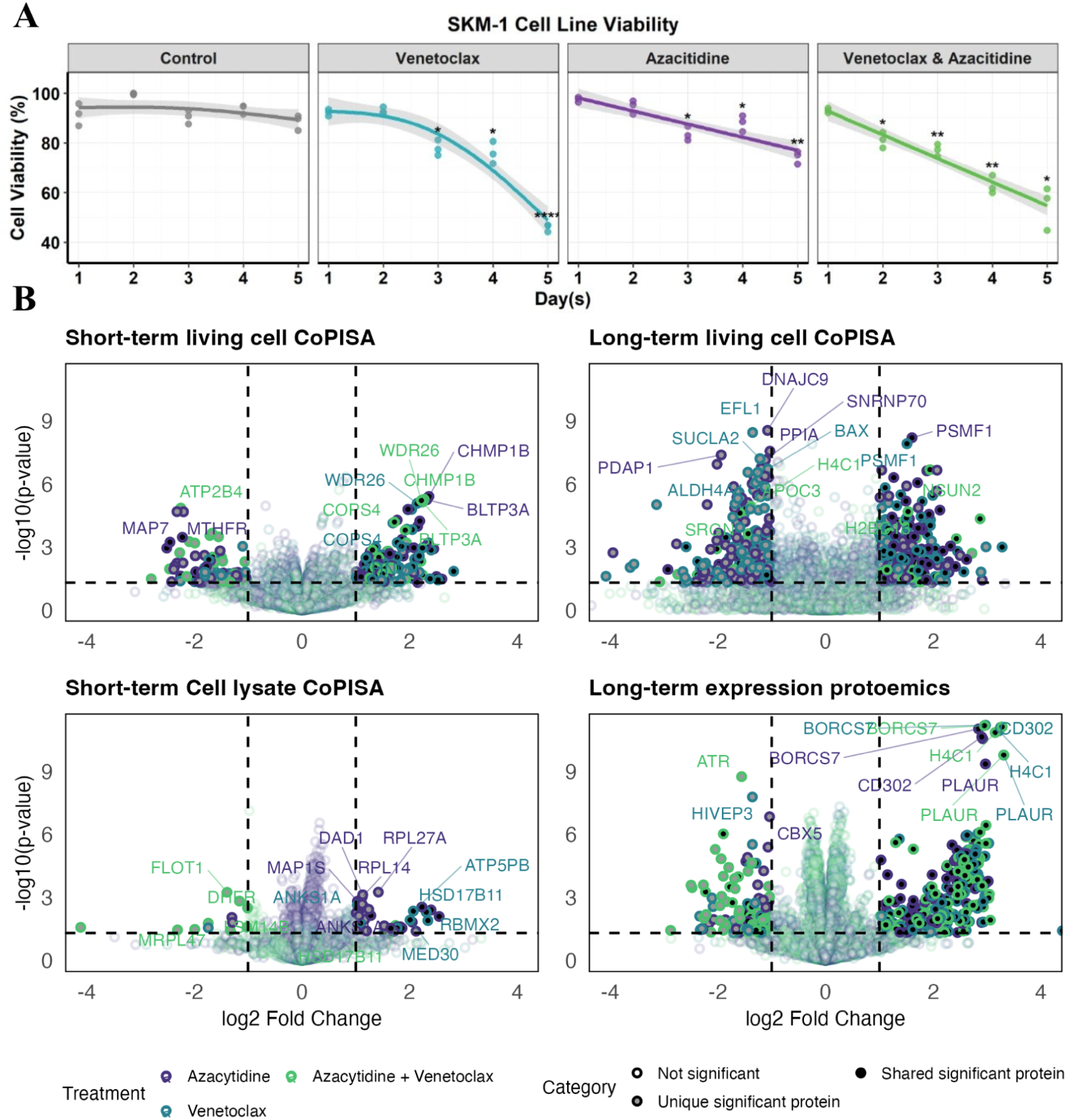
Viability-guided long-term treatment selection and multi-layer proteomic profiling. **(A)** SKM-1 cells were treated with azacitidine (300 nM), venetoclax (50 nM), venetoclax plus azacitidine, or DMSO control for 5 days. Cell viability was measured every 24 h using the trypan blue exclusion assay. Data points represent mean viability values, and error bars indicate standard deviation. Nonlinear regression curves were fitted to visualize viability trends over time. Statistical significance was calculated by comparing each time point with the first measured time point within the same treatment group, with p-values adjusted using the Benjamini–Hochberg method. Asterisks indicate significance: p < 0.05, p < 0.01, p < 0.001, p < 0.0001. **(B)** Each volcano plot overlays three treatment-versus-control comparisons, with colors indicating azacytidine (300 nM; purple), venetoclax (50 nM; turquoise), and the combination treatment VA (yellow). The plots display solubility shifts detected in short-term living-cell CoPISA, short-term cell-lysate CoPISA, and long-term living-cell CoPISA assays, as well as differential protein expression measured in the long-term expression proteomics assay. Proteins uniquely identified in a single treatment condition are shown as gray-filled dots, whereas proteins identified across multiple treatments are shown as black-filled dots. Statistical significance was evaluated using linear models with empirical Bayes moderation implemented in the limma framework. For each comparison, two-sided moderated t-tests were performed between treatment and control groups. Volcano plots show log2-transformed fold changes versus −log10-transformed P values. False discovery rates were estimated using a permutation-based approach to ensure robust significance assessment (see Methods). Source data is provided in the accompanying Source Data file.

The VA combination showed a gradual and consistent decrease in viability over time, reaching approximately 50% viability on day 5. Based on this result, day 5 was selected as a long-term treatment endpoint, representing a condition in which a substantial fraction of cells had been eliminated while a heterogeneous surviving population remained. This residual population was expected to include a mixture of drug-tolerant, slowly proliferating, and potentially still-declining cells. Nevertheless, this intermediate survival window provides a useful model to study early drug tolerance and adaptive survival responses under sustained treatment. Accordingly, surviving cells at day 5 were used for long-term CoPISA and expression proteomics to investigate candidate adaptive and resistance-associated mechanisms.

To identify proteins affected by venetoclax, azacitidine, and their combination, we compared drug-treated samples with control across four proteomic layers: short-term living-cell CoPISA (SL), short-term cell-lysate CoPISA (SC), long-term living-cell CoPISA (LL), and long-term expression proteomics (LE). Candidate proteins were selected based on fold-change and statistical criteria, and the results are summarized in the volcano plots shown in **Fig. 2B**.

In the short-term living-cell CoPISA dataset, all three treatments induced detectable protein solubility changes, indicating early drug-induced effects in intact cells. Several proteins showed strong solubility shifts, including MAP7, ATP2B4, BLTP3A, WDR26, and CHMP1B. These early changes may reflect the immediate cellular response to drug exposure, including both direct and indirect effects of treatment.

The short-term cell-lysate CoPISA dataset showed a more restricted pattern of candidate solubility changes compared with living cells. Because lysate-based CoPISA is performed after cell disruption, this module is more suitable for detecting direct or proximal protein-state changes for compounds that can engage proteins in cell extracts [15,21]. However, this interpretation does not apply equally to azacitidine, which requires cellular uptake, phosphorylation, and incorporation into RNA and DNA for its canonical activity. Therefore, the 15-min azacitidine lysate condition should not be interpreted as direct azacitidine target engagement. Instead, it should be considered an exploratory extract-level comparison that measures protein solubility changes observed after exposure to the compound under lysate assay conditions. In the VA lysate condition, detected effects may therefore reflect venetoclax-driven effects, non-canonical extract-level effects, or assay-context effects rather than canonical azacitidine biology. Proteins such as *FLOT1, SNUPN, ANKS1A, RPL14, RPL27A, ATP5PB,* and *HSD17B11* showed marked solubility changes in this layer.

In contrast, the long-term living-cell CoPISA dataset showed a broader distribution of candidate solubility changes after 5 days of treatment. This suggests that surviving cells undergo extensive adaptation under prolonged drug exposure. Several proteins, including DNMT1, DNAJC9, PSMF1, NSUN2, H4C1, and APOC3, were among the top altered proteins. The DNMT1 signal is mechanistically notable because DNMT1 is a canonical downstream target of azacitidine-associated DNA methylation biology after cellular uptake, metabolic activation, and incorporation into DNA [22]. However, the DNMT1 CoPISA signal should not be interpreted as direct drug binding; rather, in the long-term intact-cell VA condition, altered DNMT1 solubility may reflect azacitidine-associated DNMT1 trapping, chromatin association, depletion, altered complex formation, or adaptive remodeling of DNMT1-containing protein states. These long-term solubility changes may represent proteins involved in drug tolerance, persister-like adaptive states, or adaptive remodeling of the proteome.

The long-term expression proteomics dataset showed a clear shift toward increased protein abundance in many drug-treated samples. Proteins with increased abundance included PLAUR, H4C1, BOK, CD302, BORCS7, and ATR, suggesting that prolonged treatment activates compensatory survival programs. Short-term lysate-based CoPISA is expected to be more enriched for direct or proximal drug-associated protein effects, because drug exposure occurs after cell disruption. Short-term living-cell CoPISA can capture early protein solubility or stability changes occurring in intact cells, including both direct target engagement and proximal cellular responses. In contrast, long-term living-cell CoPISA reflects cumulative solubility changes in surviving cells after prolonged treatment, which may arise from direct or indirect mechanisms, including post-translational modifications, altered subcellular localization, and changes in protein–protein complex formation.

Together, these results show that venetoclax and azacitidine produce different proteomic effects depending on treatment duration and experimental context. Short-term CoPISA highlights early drug-responsive proteins, while long-term CoPISA and expression proteomics reveal adaptive changes in surviving cells. This multi-layer strategy provides a framework to identify proteins that may contribute to resistance after prolonged VA treatment, as well as novel resistance-associated drug targets.

To ensure the robustness of the significance threshold and assess the empirical false-positive rate of our analytical workflow, a comprehensive permutation analysis of protein abundance values was performed across four independent comparison datasets. For each analysis, sample labels were randomly permuted to generate 10,000 null datasets, and the same statistical testing pipeline and selection criteria applied to the original data were used to estimate the number of significant hits expected under random assignment. Across all four analyses, the resulting permutation-based global p-values were consistently low, indicating that the observed numbers of significant proteins were unlikely to have arisen by chance. The permutation-based global false discovery rates (FDRs) ranged from 2.01% to 7.82% (SL: 7.82%, LL: 5.34%, SC: 3.12%, and LE: 2.01%), demonstrating consistent deviation from the empirical null distribution across all experimental conditions.

### Intersection analysis identifies treatment-specific and shared protein responses

To better understand how venetoclax, azacitidine, and their combination affect the proteome, we next compared the overlap of candidate proteins among the three drug-treated conditions (A, V, and VA) within each proteomic layer. (**Fig. 3**). Consistent with our previous demonstration of conjunctive targeting, where drug combinations generate emergent combination-specific protein targets beyond those detected with individual agents, we examined whether VA-responsive proteins represented a simple additive combination of single-drug responses or contained additional combination-specific changes.

**Figure 3:**
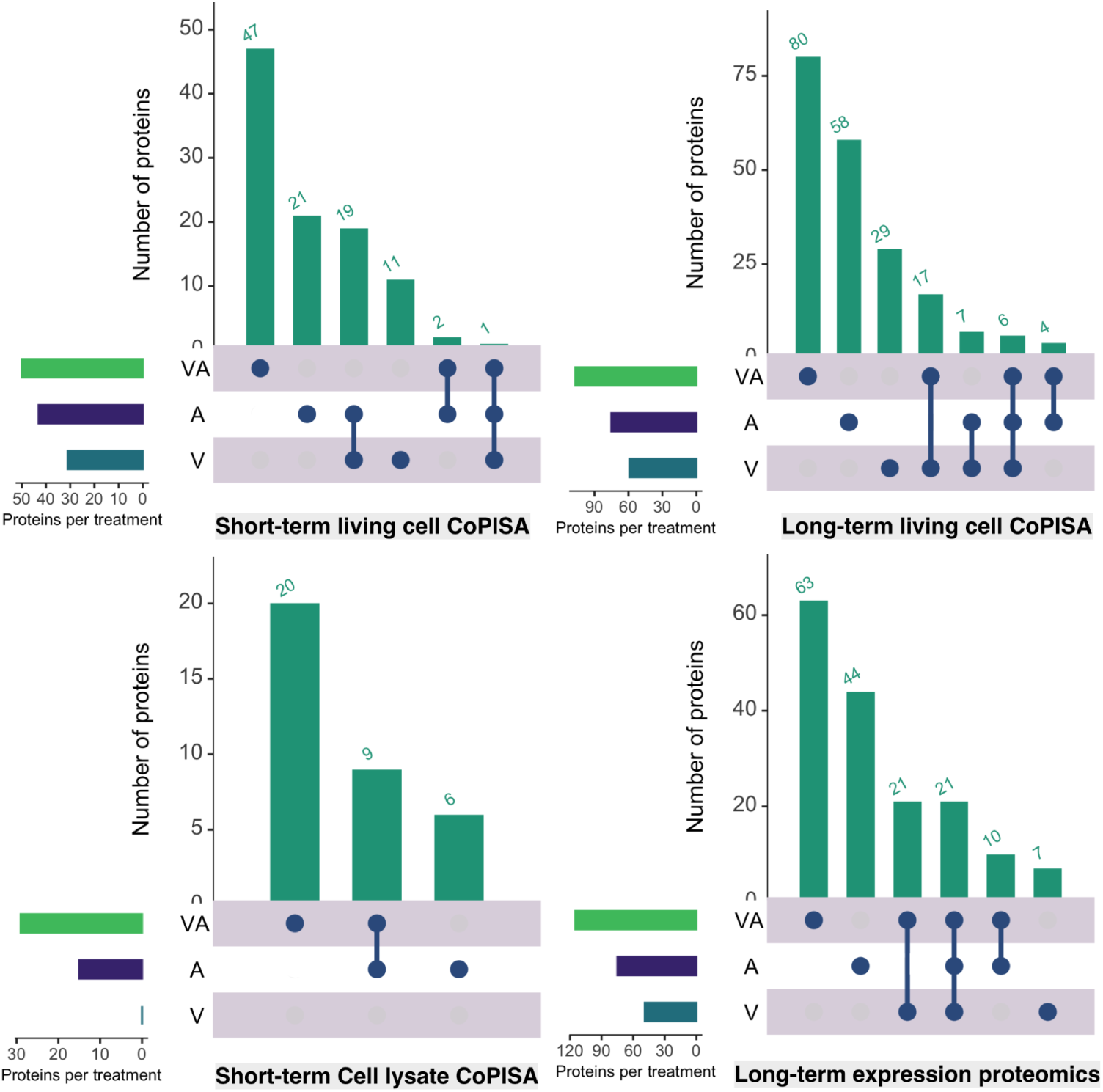
Intersection analysis of protein responses identified across four proteomic assays following single-agent and VA combination treatment. Each UpSet plot summarizes the overlap and treatment-specific distribution of candidate proteins identified after treatment with venetoclax (V), azacitidine (A), or their combination (VA; Venetoclax–Azacitidine) across the four proteomic workflows. The plots display proteins uniquely associated with each treatment condition, as well as proteins shared between single-agent and combination responses, highlighting both conserved and combination-specific proteomic changes under distinct cellular contexts. Statistical evaluation of overlap enrichment and VA-specific response fractions and source data are provided in the accompanying Source Data file.

In the short-term living-cell CoPISA dataset, the largest group of candidate proteins was specific to the VA combination, with 47 proteins detected only after combined treatment. This suggests that venetoclax and azacitidine together induce early protein solubility changes that are not observed with either single drug alone. Smaller groups of proteins were shared between treatments, including proteins common to azacitidine and venetoclax, or shared across all treatment groups. Statistical evaluation of treatment overlap using hypergeometric enrichment analysis supported that the observed overlap between VA- and azacitidine-responsive proteins was greater than expected by chance (9 shared proteins versus 0.14 expected; 65.7-fold enrichment; FDR < 0.001). However, the majority of VA-responsive proteins remained combination-specific, with 20 of 29 VA-associated proteins not detected in either single-drug response (69% VA-specific fraction). No significant overlap was detected between VA and venetoclax in this dataset, consistent with the absence of significant venetoclax-only responses in this short-term module.

In the short-term cell-lysate CoPISA dataset, fewer candidate proteins were detected overall. Lysate-based CoPISA can be useful for detecting direct or proximal protein-state changes for drugs that engage proteins in cell extracts. However, this interpretation does not apply equally to azacitidine. Azacitidine requires cellular uptake, metabolic activation through phosphorylation, and incorporation into RNA and DNA to exert its canonical biological effects [23,24]. Therefore, the 15-min lysate-based azacitidine condition should not be interpreted as direct azacitidine target engagement.

Instead, the azacitidine lysate module was included as an exploratory extract-level comparison. It measures whether exposure of the lysate to the azacitidine compound or to the venetoclax–azacitidine mixture is associated with detectable protein solubility changes under the assay conditions. These changes may reflect non-canonical extract-level effects, assay-context effects, or, in the combination condition, effects driven mainly by venetoclax rather than canonical azacitidine biology. Interpretation of azacitidine activity should therefore rely primarily on the intact-cell short-term module and the long-term treatment modules, where cellular uptake, metabolism, and downstream RNA/DNA-associated effects can occur. Despite the lower number of detected candidates in this module, VA-responsive proteins remained predominantly combination-specific, with 47 of 50 VA-associated proteins not observed in either single-drug condition (94% VA-specific fraction).

The long-term living-cell CoPISA dataset showed a broader overlap pattern, with 59 proteins specific to VA and 46 proteins specific to venetoclax. This indicates that after prolonged treatment, surviving cells retain distinct solubility changes, especially in response to the combination and venetoclax-alone modules. These proteins may reflect DTP cell states or adaptive changes that remain detectable after 5 days of treatment. Hypergeometric analysis demonstrated that the shared components between VA and individual treatments were significantly enriched above random expectation, including VA–venetoclax overlap (23 shared proteins versus 1.99 expected; 11.6-fold enrichment; FDR < 0.001) and VA–azacitidine overlap (10 shared proteins versus 2.53 expected; 4-fold enrichment; FDR < 0.001). Nevertheless, 75% of VA-responsive proteins remained unique to the combination treatment, supporting the presence of an emergent VA-specific response rather than a simple summation of single-drug effects.

In the long-term expression proteomics dataset, the largest groups were again treatment-specific, with 60 VA-specific proteins and 50 venetoclax-specific proteins. This indicates that long-term treatment strongly remodels protein abundance, particularly in cells exposed to venetoclax or the VA combination. These expression changes may represent compensatory survival mechanisms that help cells adapt to prolonged drug pressure. In this dataset, VA also displayed significant overlap with both individual treatments, with VA–azacitidine and VA–venetoclax overlaps exceeding random expectation (17-fold and 35-fold enrichment, respectively; FDR < 0.001). Importantly, despite these shared responses, 55% of VA-responsive proteins were not detected in either single-drug condition, demonstrating that the combination treatment induces additional protein abundance changes beyond those associated with venetoclax or azacitidine alone.

Collectively, these results show that the VA combination produces a distinct proteomic response across both early and long-term treatment settings. Rather than representing a completely independent response from the individual drugs, VA integrates shared drug-associated effects with additional combination-specific molecular changes. The statistically significant but partial overlap between VA and single-drug responses indicates that the combination preserves components of venetoclax and azacitidine biology, while the substantial fraction of VA-specific proteins across all datasets demonstrates that VA is not equivalent to the additive combination of individual drug responses. These findings extend the concept of conjunctive targeting by showing that rational drug combinations can generate emergent proteomic signatures that cannot be predicted solely from single-agent responses.

### Cross-layer integration reveals persistent proteomic signatures associated with long-term drug adaptation

We next compared candidate proteins across the four proteomic layers for each treatment separately. This analysis allowed us to identify proteins that were detected in short-term CoPISA datasets, long-term CoPISA datasets, and long-term expression proteomics (**Fig. 4A**). Because these workflows capture distinct aspects of protein biology, including early protein solubility changes, persistent solubility alterations, and long-term protein abundance remodeling, we further evaluated whether proteins shared between proteomic layers exceeded random expectation using pairwise hypergeometric enrichment analysis.

**Figure 4.**
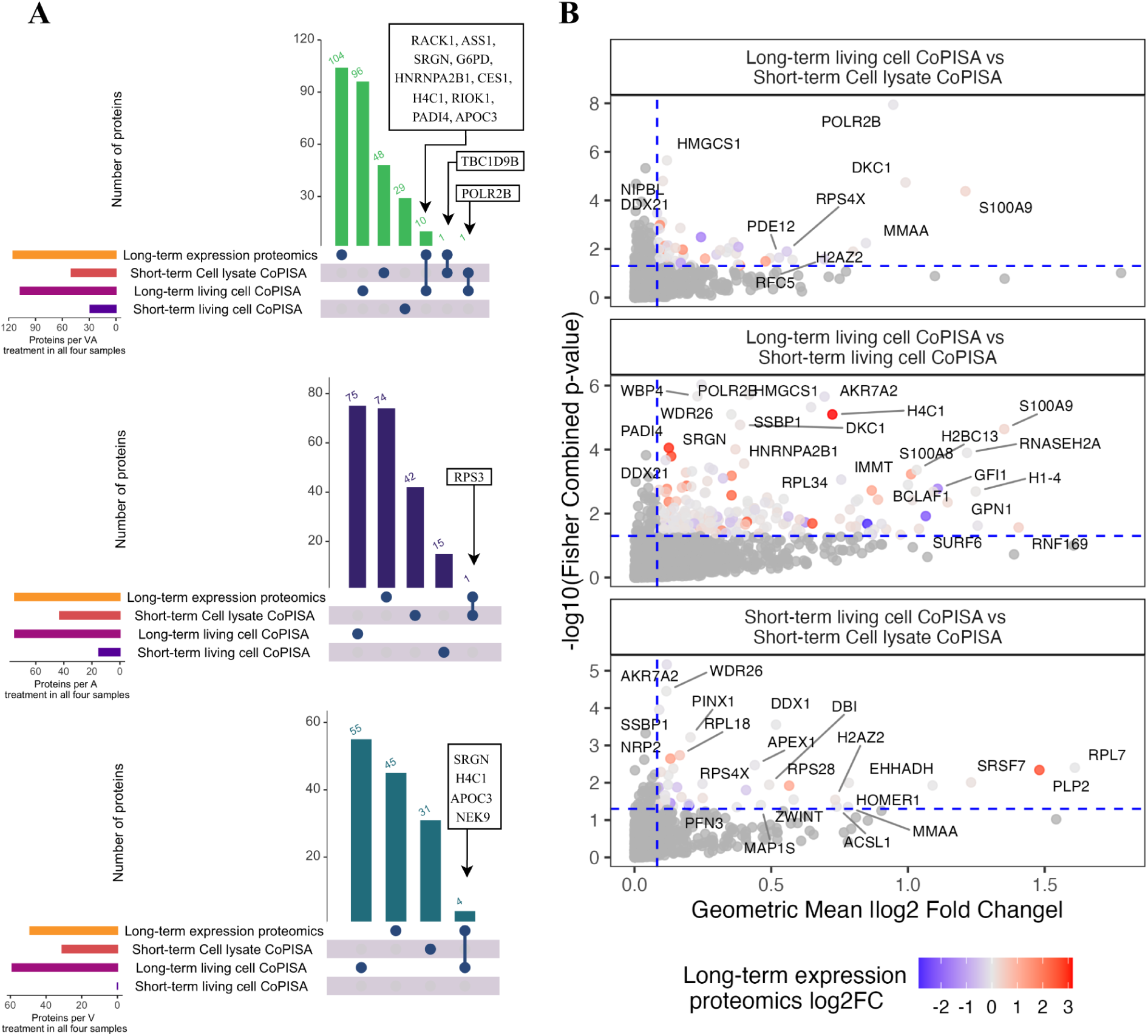
Cross-assay overlap and integrated solubility profiling of treatment-associated proteins across CoPISA and proteomic assays. **(A)** Each UpSet plot depicts the overlap of significant proteins identified under a specific treatment condition, VA (Venetoclax–Azacitidine), azacitidine (A), or venetoclax (V), across four proteomic workflows: short-term living-cell CoPISA, short-term cell-lysate CoPISA, long-term living-cell CoPISA, and long-term expression proteomics. Labeled proteins represent shared hits detected across multiple experimental layers, highlighting a subset of persistent treatment-associated molecular responses. **(B)** Pairwise comparison of proteome-wide solubility shifts in the VA treatment across CoPISA assays. Each scatter plot (spray plot) shows combined solubility effect sizes (geometric mean of absolute log₂ fold changes) derived from short-term living-cell CoPISA, short-term cell-lysate CoPISA, and long-term living-cell CoPISA, plotted against statistical significance summarized as −log₁₀(Fisher’s combined p-values). Each point represents a protein, with those passing both significance and effect-size thresholds highlighted and color-scaled according to long-term expression proteomics log₂ fold change. The effect-size cutoff corresponds to the 75th percentile of absolute geometric mean solubility changes across all proteins (see Methods). Dashed lines indicate significance (p = 0.05) and effect-size thresholds. Selected proteins are labeled for clarity. Source data is provided in the accompanying Source Data file.

For the VA combination, most proteins were specific to the long-term expression proteomics layer, indicating that prolonged combination treatment strongly changes protein abundance in surviving cells. A smaller subset of proteins was shared across multiple layers, including RACK1, ASS1, SRGN, G6PD, HNRNPA2B1, CES1, H4C1, RIOK1, PADI4, APOC3, TBC1D9B, and POLR2B (**Fig. 4A**). These proteins may be especially important because they connect early drug response with long-term adaptation. Hypergeometric analysis showed that significant convergence was observed only between the long-term living-cell CoPISA and long-term expression proteomics datasets (10 shared proteins versus 2.2 expected by chance; 4.5-fold enrichment; FDR = 3.8 × 10⁻⁴), whereas all other pairwise overlaps were not greater than expected by chance. This finding indicates that the majority of proteins identified in each proteomic layer represent orthogonal rather than redundant information, while the subset shared between the two long-term assays likely reflects persistent molecular adaptations that are detectable both as altered protein solubility and altered protein abundance.

For azacitidine, the strongest signal was also observed in long-term expression proteomics, with many proteins detected only after prolonged treatment. Only a small number of proteins overlapped across layers, including RPS3, suggesting that azacitidine alone induces a more limited shared response across the CoPISA and expression datasets. Consistent with this observation, none of the pairwise overlaps between proteomic layers reached statistical significance in the hypergeometric analysis, indicating that each proteomic workflow captured largely distinct aspects of the azacitidine response. This observation is consistent with the delayed and indirect mechanism of action of azacitidine, which requires cellular uptake, metabolic activation, and downstream incorporation into RNA and DNA before producing measurable proteomic consequences.

For venetoclax, several proteins were detected across long-term layers, including SRGN, H4C1, YBX1, and NEK9. These proteins may represent venetoclax-associated adaptive responses in surviving cells. Their presence in long-term CoPISA and/or expression proteomics suggests that they may contribute to drug tolerance or resistance after prolonged venetoclax exposure. Similar to the VA combination, significant overlap was detected only between long-term living-cell CoPISA and long-term expression proteomics (4 shared proteins versus 0.52 expected; 7.7-fold enrichment; FDR = 0.01), whereas all remaining pairwise comparisons were indistinguishable from random. These results further support the view that persistent protein-state alterations measured by long-term CoPISA are linked to corresponding changes in protein abundance during prolonged venetoclax treatment.

In general, this cross-layer comparison shows that most treatment-associated changes are specific to a single proteomic layer, particularly long-term expression proteomics. Hypergeometric analysis further demonstrated that overlap between proteomic workflows was generally limited and, with the exception of the long-term living-cell CoPISA and long-term expression proteomics datasets, did not exceed random expectation. These findings indicate that the four proteomic workflows provide largely orthogonal rather than redundant information, capturing distinct dimensions of the cellular response to therapy. The significant convergence between the two long-term assays for the VA combination and venetoclax treatment identifies a small but robust set of proteins that persist across independent proteomic measurements and are therefore strong candidates for mediating long-term adaptation and drug tolerance. In contrast, the predominance of layer-specific proteins highlights the value of integrating multiple proteomic strategies to comprehensively characterize drug-induced molecular responses that would not be captured by any single workflow alone.

### Cross-layer integration identifies proteins shared across CoPISA datasets

To identify proteins showing consistent responses across multiple proteomic layers, we integrated the CoPISA datasets using Fisher’s method to combine statistical evidence [25] and a geometric mean fold-change metric to summarize effect size (**Fig. 4B**). Long-term expression proteomics was incorporated as a color annotation layer, allowing visualization of whether proteins with shared CoPISA responses also displayed abundance changes after prolonged treatment.

The comparison between long-term living-cell CoPISA and short-term cell-lysate CoPISA identified a relatively small set of shared proteins, including POLR2B, DKC1, MMAA, and S100A9 (**Fig. 4B**). Because lysate-based CoPISA is enriched for direct or proximal drug-associated effects, proteins detected in both datasets may represent persistent biochemical responses that remain detectable after prolonged treatment. Among these proteins, S100A9 showed one of the strongest combined responses and was also associated with increased abundance in the long-term expression dataset.

The comparison between long-term living-cell CoPISA and short-term living-cell CoPISA identified a larger group of proteins with significant responses across both intact-cell datasets (**Fig. 4B**). These included H4C1, S100A9, RNASEH2A, H1-4, HLA-A, LMNB2, HNRNPA2B1, DKC1, WDR26, and AKR7A2. Several of these proteins map onto the established mechanism of action of azacitidine rather than representing a novel biological signature: the chromatin-associated histones H4C1 and H1-4, together with the nuclear structural protein LMNB2, are consistent with chromatin reorganization downstream of DNMT1 depletion and hypomethylation [22], and HNRNPA2B1, DKC1, and RNASEH2A, involved in RNA processing, rRNA pseudouridylation, and resolution of RNA:DNA hybrids, respectively, reflect the RNA metabolism disruption caused by azacitidine’s incorporation into RNA and DNA [23,24]. This convergence is therefore expected given azacitidine’s canonical pharmacology, and we interpret it as internal validation that CoPISA captures genuine, mechanistically relevant drug engagement rather than stochastic solubility noise.

Intersection of short-term living-cell CoPISA and short-term cell-lysate CoPISA identified proteins shared between the two acute-response models (**Fig. 4B**). These included APEX1, DDX1, RPS28, RPL7, PLP2, SRSF7, EHHADH, H2AZ2, HOMER1, ACSL1, MAP1S, PFN3, and DBI. Because these proteins were detected in both intact-cell and lysate contexts shortly after treatment, they may represent early VA-responsive protein states that are less dependent on long-term cellular adaptation. Notably, several proteins in this group are associated with RNA processing, ribosome function, lipid metabolism, and endoplasmic reticulum stress pathways.

Overall, the cross-layer integration analysis identified a subset of proteins that remained significant across multiple CoPISA datasets, suggesting reproducible solubility responses to VA treatment. The expression-proteomics overlay further showed that some of these proteins also undergo abundance remodeling after prolonged treatment, whereas others display strong solubility changes with limited expression changes. These findings support the idea that protein-state remodeling and protein-abundance remodeling represent complementary dimensions of the AML response to VA treatment.

### Integrated CoPISA–expression analysis prioritizes retained and remodeled adaptive targets

We next applied the integrative framework introduced above to organize treatment-responsive proteins based on their behavior across short-term CoPISA, long-term CoPISA, and long-term expression proteomics. This analysis was intended to compare proteins exhibiting early biochemical perturbations with those retaining or losing such responses under prolonged treatment in order to characterize features associated with the residual, drug-exposed state. As described in the introduction, we grouped proteins into four regimes (R1–R4) as a structured and interpretable way to summarize combined patterns across datasets: sensitive-persistent common targets (R1), persistent-adaptive dosage-compensation targets (R2), sensitive-specific targets (R3), and adaptive-remodeled targets (R4). Rather than constituting a definitive biological taxonomy, these regimes provide a heuristic stratification to facilitate interpretation of coordinated solubility and abundance changes across conditions. Within this framework, R2 and R4 represent protein groups of particular interest in the context of treatment adaptation, as they combine drug-associated changes in solubility behavior with increased abundance in the long-term surviving population, suggesting potential roles in adaptive or compensatory responses.

In **Fig. 5A**, the right panel represents the retained-target branch of this classification. Proteins in this group were significant in short-term CoPISA and remained significant in long-term living-cell CoPISA, indicating that their solubility or stability alteration persisted in the surviving cell population after prolonged VA exposure. Retained proteins with limited long-term abundance increase correspond to the sensitive–persistent common target class, R1, suggesting sustained biochemical responsiveness without clear evidence of compensatory expression. R1 and R2 proteins showed significant enrichment for RNA-processing and spliceosome-associated processes, including RNA splicing via transesterification reactions, mRNA splicing, and mRNA processing. These terms were driven by *SYNCRIP, HNRNPR, SNRPE, HNRNPA1, HNRNPA1L3, and HNRNPA1L2*, indicating that retained CoPISA targets are strongly connected to post-transcriptional RNA regulation (**Fig. 6A & Supplementary File S1**). This is biologically relevant because RNA splicing and mRNA processing can reshape protein isoform usage, apoptotic signaling, and therapy response in cancer [26]. In leukemia, modulation of RNA splicing has also been linked to altered response to BCL2 inhibition [27], supporting the interpretation that persistent solubility changes in RNA-processing proteins may reflect a sustained adaptive layer of VA response.

**Figure 5:**
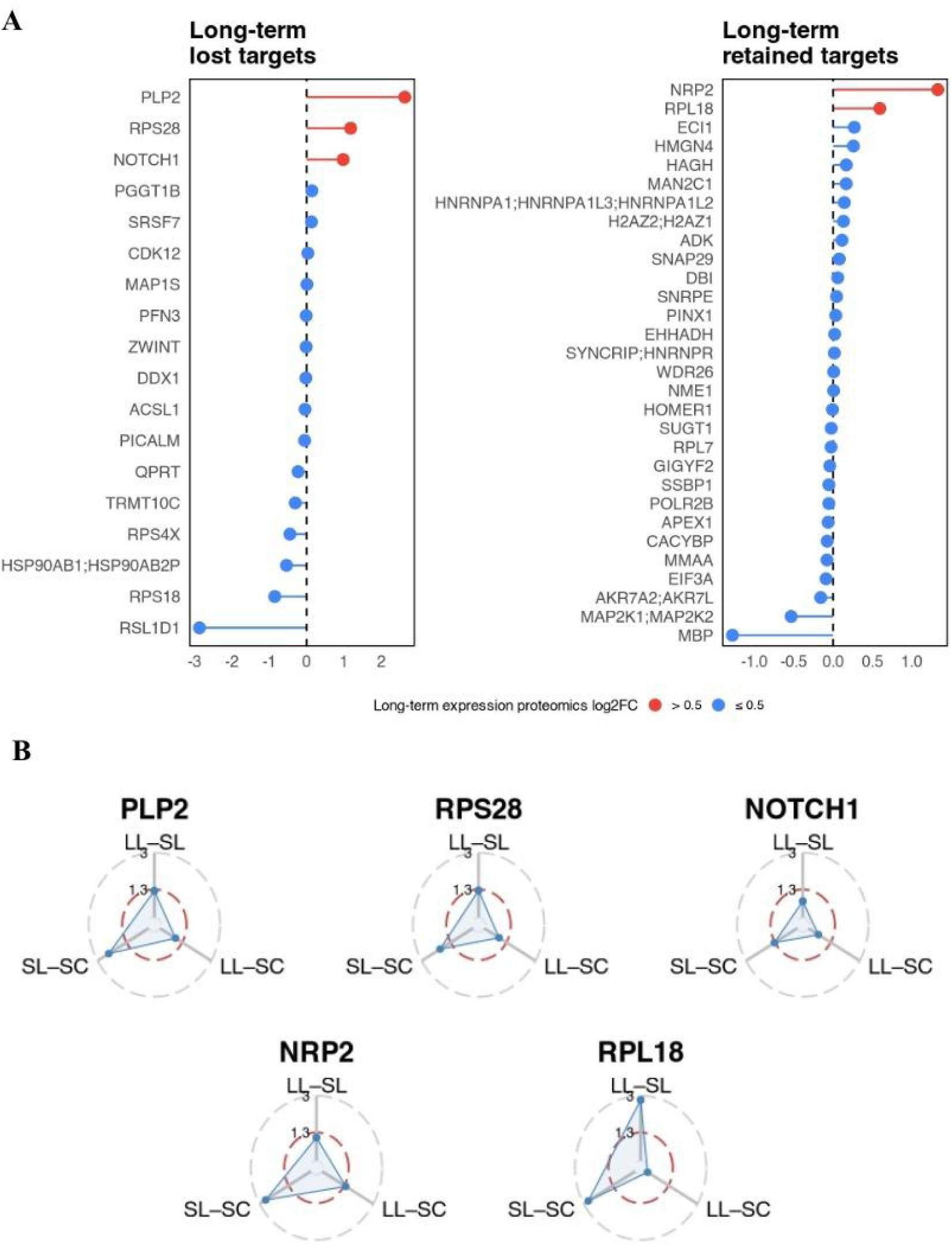
Long-term lost and retained protein targets based on short- and long-term CoPISA and expression proteomics. **(A)** The left panel shows proteins classified as “long-term lost targets,” corresponding to short-term CoPISA hits that are not retained as significant in long-term CoPISA (R3 and R4). The right panel shows “long-term retained targets,” representing proteins that remain significant in long-term CoPISA (R1 and R2). Within each category, proteins are ranked by long-term expression proteomics log₂ fold change, with color indicating magnitude thresholds (>0.5 vs ≤0.5). **(B)** Radar plots show the −log₁₀-transformed Fisher combined p-values for the three pairwise comparisons among long-term living-cell CoPISA (LL), short-term living-cell CoPISA (SL), and short-term cell-lysate CoPISA (SC) for selected proteins with high abundance changes in long-term expression proteomics. The dashed red reference line indicates the significance threshold (p = 0.05; −log₁₀(p) = 1.3), with values extending beyond the threshold indicating significant combined evidence. Source data is provided in the accompanying Source Data file.

**Figure 6.**
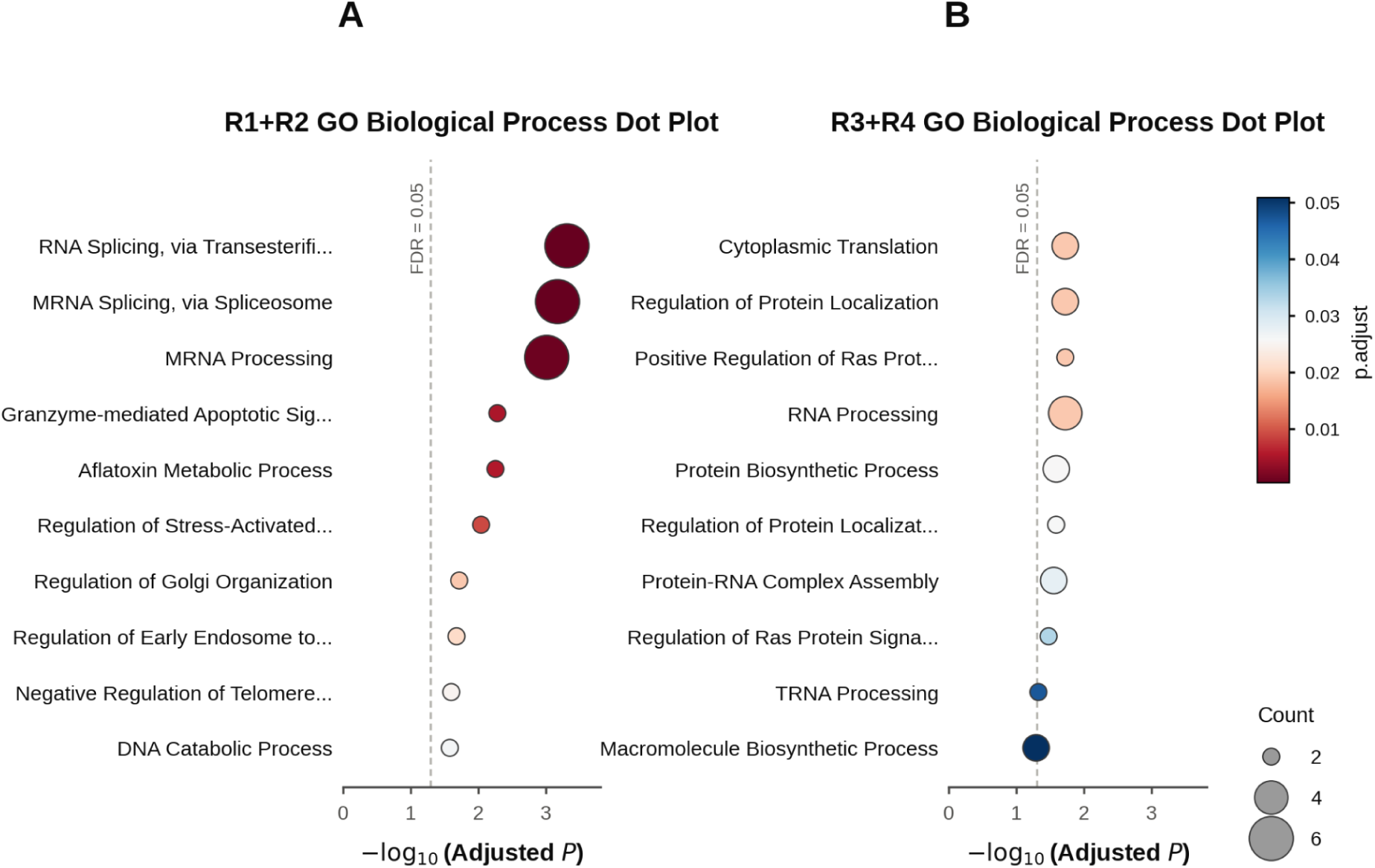
GO Biological Process enrichment analysis of retained (R1 & R2) and lost (R3 & R4) CoPISA target proteins. (A) Top 10 enriched GO Biological Process terms among R1 & R2 proteins (sensitive–persistent common targets: short-term CoPISA hits that remained significant in long-term living-cell CoPISA without strong long-term expression induction). (B) Top 10 enriched terms among R3 & R4 proteins (sensitive-specific targets: short-term CoPISA hits not retained as significant in long-term CoPISA). Gene set enrichment was performed using Enrichr against the GO Biological Process gene set library. Dot color indicates the Benjamini–Hochberg-adjusted P-value (p.adjust) and dot size indicates the number of proteins annotated to each term (Count); both scales are shared across panels A and B to allow direct comparison of enrichment magnitude between the retained and lost target classes. Terms are ranked by −log₁₀(adjusted P-value); the dashed line marks the FDR = 0.05 significance threshold. Full enrichment results for all tested terms are provided in Supplementary File S1.

R1 and R2 were also enriched for stress-activated kinase signaling, the ERK1/ERK2 cascade, and regulation of protein serine/threonine kinase activity, all driven by the same two genes, MAP2K1 and MAP2K2 **(Fig. 6A & Supplementary File S1)**. Because RAS/MAPK pathway activation can promote MCL1-mediated venetoclax resistance in AML [28], persistent biochemical alteration of MAPK-related proteins may represent a survival-associated signaling module in the long-term treated population. Two additional significant terms, regulation of Golgi organization and regulation of early endosome-to-late endosome transport, were also annotated to MAP2K1 and MAP2K2 rather than reflecting independent vesicle-trafficking evidence. Beyond this MAPK-linked cluster, the retained (R1 and R2) target set was also significantly enriched for negative regulation of telomere maintenance and of DNA biosynthetic process, driven by PINX1 and HNRNPA1, and for granzyme-mediated apoptotic signaling and DNA catabolic process, driven by APEX1 and NME1 (**Fig. 6A & Supplementary File S1**). Together, these results suggest that the retained (R1 and R2) target set is not simply a collection of passive protein hits but is organized around RNA metabolism, MAPK/ERK kinase signaling, telomere-maintenance regulation, and apoptotic/DNA-repair-associated biology. Because R1 proteins, by definition, remain significant in long-term CoPISA without a corresponding increase in long-term abundance, R1 likely captures persistent biochemical remodeling rather than dosage-compensation-driven resistance, in contrast to the abundance-linked R2 subset discussed next.

*NRP2* is the strongest R2 candidate (**Fig. 5A & B**). As a cell-surface co-receptor involved in VEGF and semaphorin signaling, migration, survival signaling, and cancer-cell plasticity, NRP2 may mark a persister-like state in which surviving AML cells engage extracellular signaling and adaptive survival programs. RPL18, a component of the 60S ribosomal subunit, links the R2 class to translational adaptation. This is relevant because altered translation, mitochondrial translation, and integrated stress-response biology have been implicated in venetoclax resistance and re-sensitization strategies in AML [28]. Therefore, the R2 pattern suggests that persistent biochemical target engagement together with abundance-level compensation may identify proteins required for maintaining survival under continued drug pressure.

To further support the prioritization of NRP2 as a retained candidate with increased abundance in R2, we examined the peptide-level MS/MS evidence for the top-ranked NRP2 peptide, GGDSITAVEAR. This peptide was consistently identified across the orthogonal proteomic modalities, including short-term living-cell CoPISA, long-term living-cell CoPISA, short-term cell-lysate CoPISA, and long-term expression proteomics. In each assay, the selected tandem mass spectrum showed reproducible fragment-ion annotation, with matched b- and y-ion series supporting confident peptide assignment. The precursor ion was detected at m/z 690.3752 with a 2+ charge state, corresponding to the 11-amino-acid NRP2 peptide GGDSITAVEAR (**Fig. 7**). The fragmentation spectra showed a consistent pattern across assays, including prominent y-ion signals and comparable annotated fragment coverage, supporting robust identification of the NRP2 peptide rather than assay-specific or low-confidence detection. The peptide identification scores were high across all modalities, including short-term living-cell CoPISA, long-term living-cell CoPISA, short-term cell-lysate CoPISA, and long-term expression proteomics. Thus, the MS/MS evidence provides peptide-level support for the presence of NRP2 across the datasets and strengthens confidence in its prioritization as a persistent-adaptive R2 candidate. While these spectra do not independently establish the functional role of NRP2 in resistance, they support the reliability of NRP2 detection in the integrated CoPISA–expression analysis.

**Figure 7:**
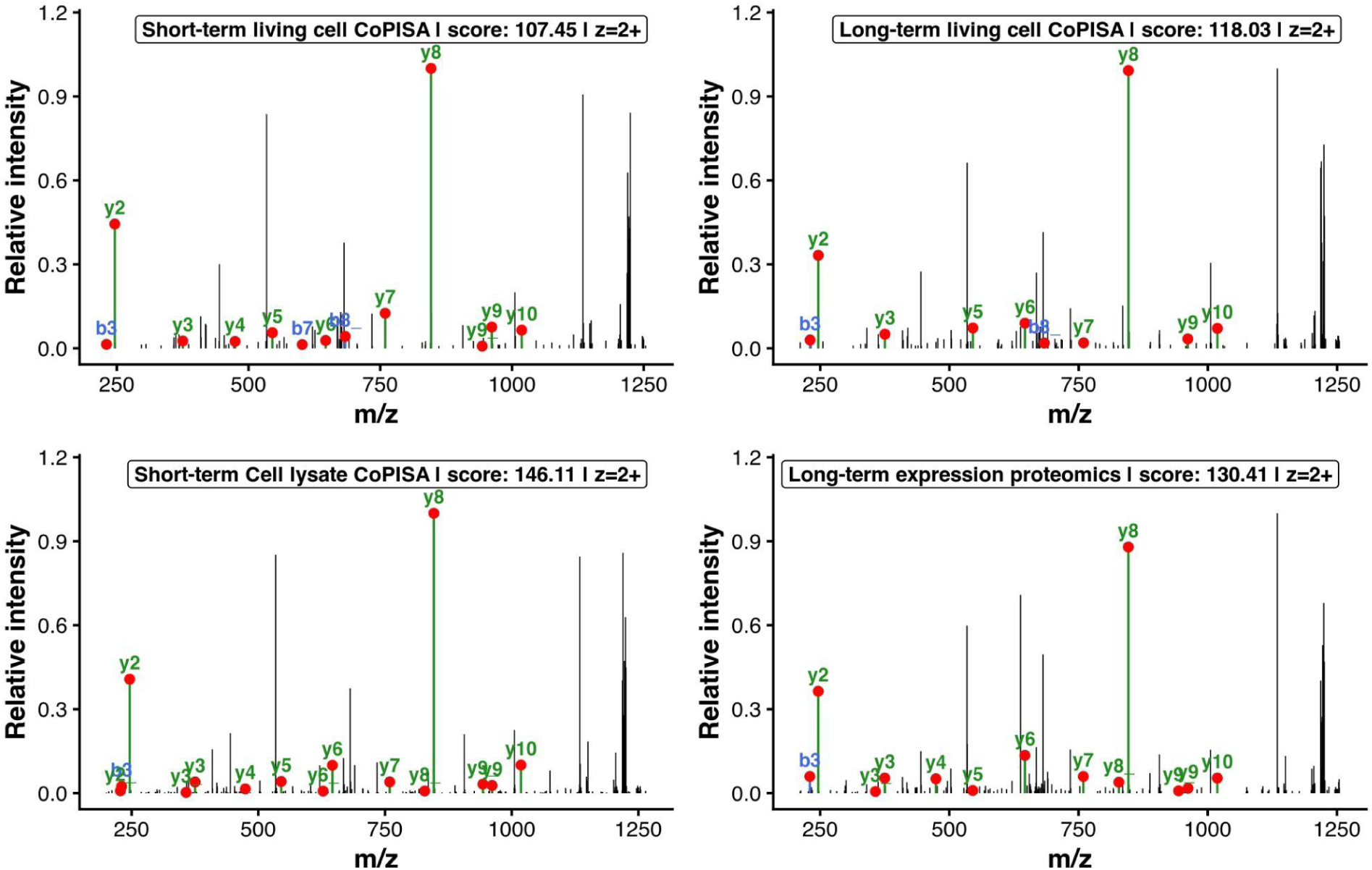
MS/MS fragmentation spectra of the top-ranked NRP2 peptide (GGDSITAVEAR) identified across proteomic modalities. Tandem mass spectra corresponding to the highest-scoring peptide derived from protein Neuropilin-2 (NRP2) are shown for each treatment. The peptide was selected based on a composite ranking score integrating spectral count, mean identification score, treatment consistency, and peptide-level confidence metrics. For each treatment, the highest-scoring MS/MS spectrum was retained and annotated against theoretical fragment ions generated from the peptide sequence. Matched fragment ions are displayed as b-ions (blue) and y-ions (green), with observed fragment peaks indicated by red markers and ion labels. Peak intensities were normalized within each spectrum to the base peak to enable comparison across conditions. Panel annotations report the treatment condition, spectrum identification score, and precursor charge state for each selected spectrum. The selected precursor had an m/z of 690.3752 (2+), corresponding to an 11-amino-acid peptide, and fragment ion assignments were performed with a tolerance of ±0.5 Da.

To further evaluate the structural context of the identified peptide, we mapped GGDSITAVEAR to the annotated NRP2 sequence and AlphaFold-predicted structure. The peptide corresponds to residues ∼507–517 within the extracellular second F5/8 type C (FA58C2) domain of NRP2 (residues 434–592). This region is predicted to adopt a well-structured conformation and is distinct from the annotated disordered regions of NRP2 (residues 298–317 and 601–622). Consistent with the AlphaFold prediction, experimentally resolved structures encompassing the extracellular region of NRP2 (residues 275–595) support a stable folded architecture. Although GGDSITAVEAR is not annotated as an individual ligand-binding motif, its localization within the FA58C2 domain places it within a functionally relevant extracellular region involved in VEGF and heparin binding. These observations provide additional support that the detected peptide originates from a structurally defined and functionally relevant region of NRP2 rather than a flexible or poorly characterized segment.

In **Fig. 5A**, the left panel represents the long-term lost-target branch of the framework. These proteins were detected as short-term CoPISA hits but were no longer significant in the long term. living-cell CoPISA, indicating that the early treatment-associated solubility response was not maintained after prolonged exposure. Lost targets with little or no increase in long-term expression correspond to sensitive-specific targets, R3, suggesting proteins that are primarily associated with the initial drug-sensitive state and are not maintained during adaptation.

To increase statistical power, GO Biological Process enrichment was instead performed on the pooled set of long-term lost targets (R3 and R4 combined; **Fig. 6B**), analogous to the pooled retained-target analysis (R1 and R2; **Fig. 6A**). This combined analysis identified nine GO Biological Process terms passing FDR < 0.05. Significant terms clustered into three themes: translation and ribosome biology (cytoplasmic translation, protein biosynthetic process, protein–RNA complex assembly), driven by the ribosomal proteins RPS4X, RPS28, and RPS18; RNA and tRNA processing (RNA processing, tRNA processing), driven by TRMT10C, CDK12, SRSF7, and DDX1; and regulation of Ras protein signal transduction and protein localization, driven by NOTCH1, PICALM, HSP90AB1, and RSL1D1 (**Fig. 6B & Supplementary File S1**). Notably, these themes directly reinforce the NOTCH1-linked signaling rewiring and RPS28-linked translational remodeling proteins discussed below as candidate R4 adaptation mechanisms. Proteins such as HMGN4, ZWINT, PINX1, MAP1S, and ACSL1, which did not contribute to any significant term, nonetheless illustrate that a subset of short-term CoPISA-responsive proteins is not maintained as long-term targets and does not undergo major abundance-level compensation, consistent with sensitive-state-associated targets that are prominent during the initial response but not retained after selection of the surviving drug-tolerant population.

By contrast, lost targets with increased long-term expression correspond to adaptive remodeled targets. *PLP2, RPS28, and NOTCH1* were among the strongest examples of this class, showing increased expression despite loss of long-term CoPISA retention. These proteins were significant in short-term CoPISA but were not retained as significant long-term CoPISA targets, despite showing increased abundance in the long-term expression proteomics layer. This lost-but-increased-in-abundance behavior is consistent with adaptive target remodeling rather than simple persistent target engagement. *PLP2* was the strongest R4 candidate and showed the largest long-term expression increase. *PLP2* is an endoplasmic-reticulum-associated membrane protein, and previous AML work has linked reduced PLP2 to ER-stress-related apoptosis and increased drug sensitivity. Therefore, increased *PLP2* abundance in the long-term surviving population may represent an adaptive mechanism that buffers ER stress or reduces apoptosis sensitivity. *RPS28*, a 40S ribosomal protein, suggests that translational remodeling is also present in the R4 class, but in this case the short-term CoPISA solubility phenotype is lost during adaptation. *NOTCH1* is especially notable because Notch signaling has been linked to AML cell survival, stromal protection, and chemoresistance. Its R4 behavior suggests that *NOTCH1* may not remain a persistent CoPISA target after prolonged treatment, but its increased abundance may reflect signaling rewiring during transition into the drug-tolerant state.

This classification therefore provides a mechanistic interpretation of the **Fig. 6A** ranking beyond a simple retained-versus-lost comparison. Proteins in the retained-and-increased-in-abundance R2 class, such as NRP2 and RPL18, may represent persistent adaptive nodes whose altered solubility and increased abundance together support survival under sustained VA pressure. In contrast, proteins in the lost-but-increased-in-abundance R4 class, such as *PLP2, RPS28, and NOTCH1*, may represent remodeled resistance-associated nodes whose early drug-responsive solubility signature is replaced by abundance-level adaptation in the long-term surviving population. Taken together, these results support the central concept that resistance-associated adaptation is not uniform but consists of separable proteomic regimes involving persistent biochemical engagement, dosage compensation, sensitive-state target loss, and adaptive target remodeling.

## Discussion

In this study, we applied a multi-layer CoPISA and expression-proteomics strategy to investigate how AML cells respond and adapt to venetoclax, azacitidine, and their combination. By integrating short-term intact-cell CoPISA, short-term lysate-based CoPISA, long-term intact-cell CoPISA, and long-term expression proteomics, we separated early drug-associated solubility changes from longer-term adaptive remodeling in surviving cells. This design is particularly relevant for VA-treated AML, where treatment failure may arise not only through fixed genetic resistance but also through transient drug-tolerant persister-like states [29,30]. Our results show that venetoclax and azacitidine induce both treatment-specific and combination-specific proteomic responses and that long-term surviving cells display distinct solubility and abundance changes that may reflect adaptive survival programs consistent with adaptive survival programs.

Because azacitidine requires cellular uptake, phosphorylation, and incorporation into RNA and DNA for its canonical mechanism of action [31], the 15-min lysate exposure was not designed to capture direct azacitidine engagement. Instead, azacitidine-containing lysate conditions were retained as exploratory extract-level controls to assess whether the VA mixture produces detectable solubility changes under lysate assay conditions. Azacitidine-related findings from this module were therefore interpreted cautiously and were not considered evidence of canonical azacitidine target engagement.

A key observation of this study is that the VA combination produced a proteomic response that was not simply the sum of the two single agents, consistent with a pattern of conjunctive targeting [12]. Across the short-term and long-term CoPISA layers, as well as in the long-term expression dataset, many candidate proteins were specific to combination treatment. This conjunctive response suggests that venetoclax and azacitidine together induce a unique cellular state involving altered protein solubility, stability, and abundance [12]. This finding is consistent with the clinical activity of venetoclax plus azacitidine in AML [1], but also with the observation that relapse after venetoclax-based regimens remains common and clinically difficult to treat [2]. Therefore, these findings highlight that the proteomic state of surviving cells after combination therapy may expose vulnerabilities that are not evident from single-agent profiling.

The comparison between short-term lysate CoPISA and short-term intact-cell CoPISA provides insight into different layers of drug action. Lysate-based CoPISA is expected to be more enriched for direct or proximal drug-associated protein effects because drug exposure occurs after cell disruption, reducing the contribution of upstream signaling, metabolism, and transcriptional adaptation [19,32]. In contrast, intact-cell CoPISA captures protein solubility changes in the cellular context and may therefore include direct target engagement, proximal pathway effects, changes in protein complexes, subcellular localization, or early stress responses [20]. The broader response observed in short-term intact cells compared with lysates supports the view that venetoclax and azacitidine rapidly perturb cellular protein networks beyond isolated drug–protein interactions.

Long-term treatment revealed a different pattern. After 5 days of VA exposure, surviving cells displayed widespread solubility and abundance changes. This indicates that the residual population is not simply an unchanged drug-insensitive background but rather represents a remodeled cellular state. This is consistent with the concept of drug-tolerant persister cells, which survive treatment through reversible and non-genetic adaptations involving altered apoptotic priming, metabolic rewiring, stress responses, epigenetic regulation, and changes in cell identity. In AML, persister-like survival has been linked to oxidative metabolism and altered physical properties of the plasma membrane [6,9,33]. Our findings extend this concept by suggesting that VA survival is also associated with extensive remodeling of protein solubility and protein abundance.

The weak correlation between CoPISA solubility changes and long-term expression changes showed that these assays capture orthogonal information. Protein abundance alone did not explain the solubility changes detected by CoPISA. This suggests that a major part of the VA response may occur at the level of protein state, including stability, complex formation, localization, or post-translational modification, rather than only through altered protein expression [21]. Cross-layer analysis identified proteins that may connect early response to long-term adaptation. In the VA condition, proteins such as RACK1, ASS1, SRGN, G6PD, HNRNPA2B1, CES1, H4C1, RIOK1, PADI4, APOC3, TBC1D9B, and POLR2B were detected across multiple proteomic layers. These proteins point toward several adaptive themes, including RNA metabolism, metabolic rewiring, chromatin regulation, ribosome-related processes, and membrane or vesicle-associated remodeling. For example, G6PD suggests involvement of the pentose phosphate pathway and redox control, while ASS1 links the response to arginine metabolism, a pathway previously explored as an AML vulnerability [34].

The integrative retained-versus-lost target framework provided a mechanistic way to interpret long-term adaptation. Retained targets without strong expression induction were enriched for RNA processing, splicing, ribonucleoprotein biogenesis, vesicle transport, MAPK signaling, and DNA-repair-related processes. RNA-processing proteins such as SYNCRIP, HNRNPR, SNRPE, and HNRNPA1-family proteins may be relevant because RNA splicing can influence leukemia response to BCL2 inhibition [27], and spliceosome biology has been associated with response to hypomethylating agent plus venetoclax therapy in AML [35]. Similarly, retained MAPK-related proteins, including MAP2K1 and MAP2K2, may be important because RAS/MAPK activation can drive MCL1-mediated venetoclax resistance in AML [36]. Because these retained targets are, by definition, the proteins whose CoPISA signal persists specifically under the combined VA condition, they correspond to the conjunctive targeting relationship defined in **Fig. 8** i.e., the logical AND-gate class [12], engaged jointly by venetoclax and azacitidine rather than by either agent alone. In **Fig. 8**, we integrate this retained/lost framework with the core VA signaling mechanism, mapping the AND-gate-classified retained proteins (R1 and R2) alongside the candidate vulnerabilities that emerge from the lost-but-increased class (R4 and R3).

**Figure 8:**
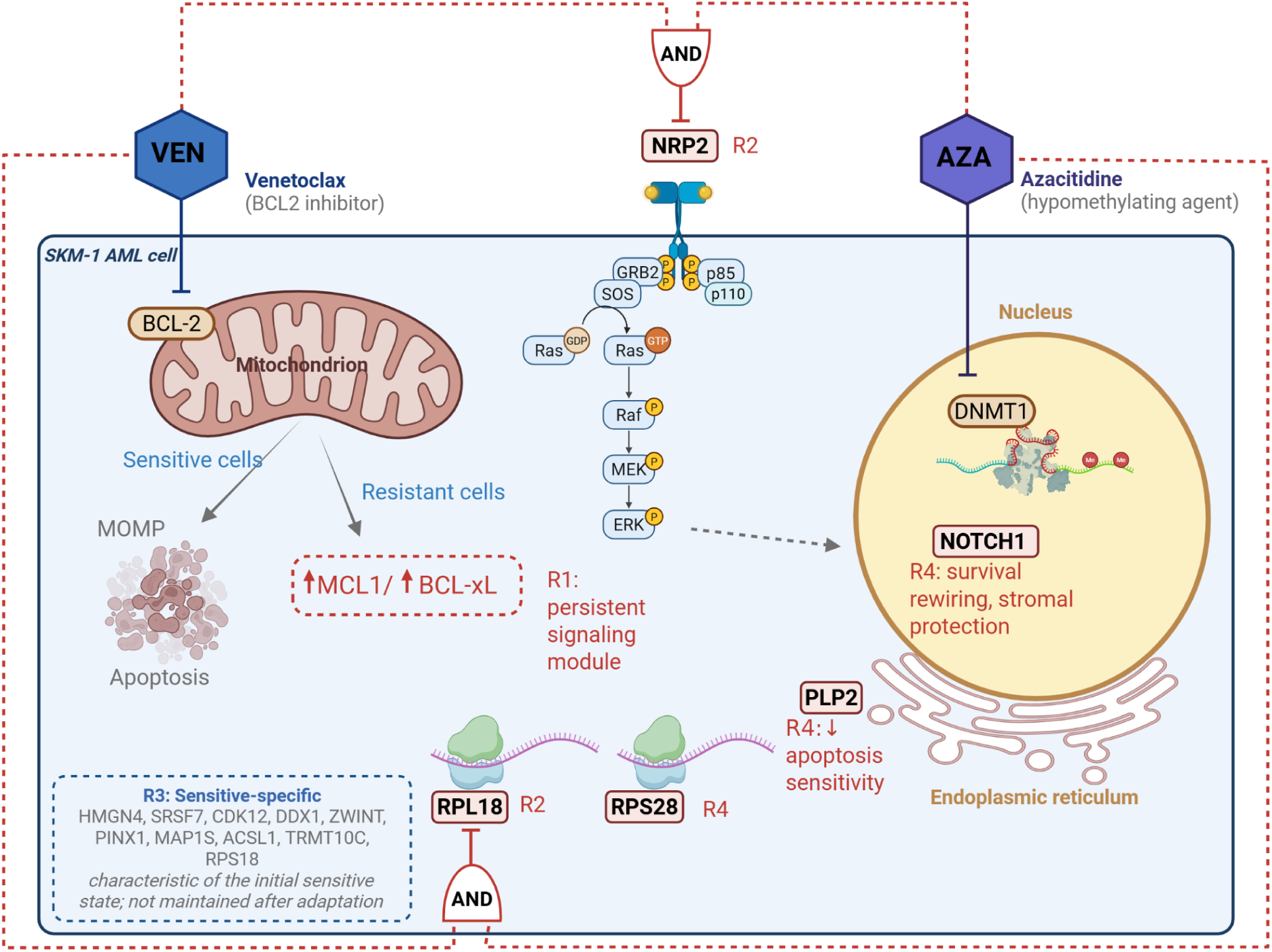
Coherent signaling model of venetoclax + azacitidine (VA) response and adaptive resistance in AML, integrating drug mechanism with CoPISA-derived proteomic classes. Coherent signaling model of venetoclax + azacitidine (VA) response and adaptive resistance in AML, with CoPISA logic-gate classification. Venetoclax inhibits BCL2, triggering BAX/BAK-mediated apoptosis in sensitive cells; azacitidine inhibits DNMT1, promoting DNA hypomethylation. Following the CoPISA logic-gate framework [12], proteins whose solubility signal is retained under combined VEN+AZA treatment (R1: RAS–MAP2K1/2–MAPK1, spliceosome/RNP granule components, H4C1/H1-4; R2: NRP2, RPL18) correspond to conjunctive targeting; the AND-gate class, jointly engaged by both drugs. Proteins whose signal is instead lost under combination but whose abundance increases long-term (R4: PLP2, RPS28, NOTCH1), together with R2 and R1, are highlighted as candidate therapeutic vulnerabilities requiring functional validation. VEN, venetoclax; AZA, azacitidine.

Among retained proteins with increased abundance, NRP2 and RPL18 emerged as key candidates. NRP2 was the strongest retained candidate with increased abundance and was supported by peptide-level MS/MS evidence across proteomic modalities. NRP2 has been linked to cancer-cell survival, migration, VEGF/semaphorin signaling, and therapy resistance in other cancer contexts [37]. However, because direct evidence for NRP2 in venetoclax-resistant AML is limited, NRP2 should be considered a candidate marker or mediator of the surviving state that requires functional validation. RPL18 links this adaptive class to ribosome biology and translation. This is relevant because mitochondrial translation and translation-associated signaling have been implicated in venetoclax resistance and resensitization in AML [38,39].

The lost-but-increased-in-abundance class highlighted a different adaptive pattern. Proteins such as PLP2, RPS28, and NOTCH1 were significant in short-term CoPISA, lost long-term CoPISA significance, but showed elevated abundance after prolonged treatment. This pattern may reflect adaptive remodeling in which the early solubility phenotype is replaced by abundance-level compensation. PLP2 is particularly interesting because reduced PLP2 has been reported to promote ER-stress-related apoptosis and increased drug sensitivity in AML [40]. Therefore, increased PLP2 levels in surviving cells may reflect ER-stress buffering. NOTCH1 may also indicate adaptive signaling rewiring, as Notch signaling has been linked to stromal-mediated AML chemoresistance, although its role in AML is context-dependent [41,42].

Overall, our results support a model in which VA adaptation is not driven by a single resistance mechanism. Instead, surviving AML cells appear to engage multiple layers of proteomic remodeling, including persistent biochemical changes, expression-level compensation, loss of sensitive-state responses, and adaptive target remodeling. This is consistent with current models of drug-tolerant persister cells, where survival is flexible, reversible, and context-dependent rather than defined by one universal marker [7,30,43].

This study has several limitations. First, the current analysis was performed in the SKM-1 AML cell line, and the findings need validation in additional AML models and primary patient samples [44]. Second, CoPISA detects changes in protein solubility or stability but does not alone prove direct drug binding. Third, the 5-day surviving population may contain a mixture of tolerant, adapting, and recovering cells. Functional experiments will be required to determine whether candidates such as NRP2, PLP2, NOTCH1, RPL18, and RPS28 are true mediators of VA tolerance or markers of the surviving state.

In conclusion, this study shows that multi-layer CoPISA combined with expression proteomics can resolve distinct proteomic regimes of VA response and adaptation in AML cells. By integrating short-term and long-term solubility changes with abundance remodeling, we identified candidate adaptive nodes, including NRP2, RPL18, PLP2, RPS28, and NOTCH1. This framework provides a systematic route to prioritize resistance-associated proteins and identify therapeutic vulnerabilities in VA-treated AML.

## Methods

### Cell culture and experimental design

SKM-1 cell line was obtained from Deutsche Sammlung von Mikroorganismen und Zellkulturen (DSMZ, Germany) and cultured in RPMI-1640 medium (Gibco, Thermo Fisher Scientific, USA) supplemented with 20% fetal bovine serum (FBS) (Gibco, Thermo Fisher Scientific), 2 mM L-glutamine (Lonza), and 100 units/mL penicillin/streptomycin (Gibco, Thermo Fisher Scientific) at 37°C and 5% CO_2_. Cells were harvested by centrifugation at 400 × g for 4 min. The cell line was tested for mycoplasma contamination using PCR-based assays and confirmed to be negative. Cells were cultured until they reached the desired density of 2 × 10⁶ cells per condition. Four treatment groups were prepared in triplicate: venetoclax (V), azacitidine (A), venetoclax plus azacitidine (VA), and DMSO-only control (C). Venetoclax was used at a final concentration of 50 nM and azacitidine at 300 nM, based on prior optimization experiments [14].

### Short-term treatments

#### Short-term (1 h) intact-cell CoPISA treatment

For short-term intact-cell profiling, SKM-1 cells were collected, counted, and adjusted to 2×10⁶ cells/mL per condition. Cells were treated with venetoclax, azacitidine, venetoclax plus azacitidine, or DMSO control in 6-well plates, with three technical replicates per condition. Cells were incubated at 37°C and 5% CO₂ for 60 min. After treatment, cells were collected into 1.5 mL tubes and pelleted by centrifugation at 600 × g for 5 min. Cell pellets were washed with PBS and centrifuged again at 600 × g for 5 min at 5°C.

Pellets were resuspended in 320 µL ice-cold PBS supplemented with protease inhibitors. Each sample was distributed into twelve 25 µL aliquots in PCR plates for thermal treatment. Aliquots were heated for 3 min across a temperature gradient from 48°C to 59°C, with 1°C increments. After heating, samples were equilibrated at room temperature for 3 min and snap-frozen in liquid nitrogen. Cells were lysed by four freeze–thaw cycles using liquid nitrogen and a 35°C heating block. Temperature-point aliquots from the same treatment and replicate were pooled into ultracentrifugation tubes. Soluble protein fractions were isolated by ultracentrifugation at 100,000 g for 20 min at 4°C. Approximately 80% of the supernatant, corresponding to ∼220 µL, was carefully collected without disturbing the pellet and used for downstream proteomics.

#### Short-term (15 min) lysate-based CoPISA treatment

For extract-based profiling, a minimum of 2×10⁶ SKM-1 cells were collected per condition. Cells were washed twice with 1× PBS to remove culture medium and pelleted at 600 × g for 5 min at 4°C. Pellets were resuspended in ice-cold PBS supplemented with protease inhibitors and lysed by four freeze–thaw cycles using liquid nitrogen and a 35°C heating block, with brief vortexing after each thaw. Lysates were clarified by centrifugation at 10,000 g for 10 min at 4°C, and clarified supernatants were recovered for drug treatment.

Clarified lysates were treated with venetoclax, azacitidine, venetoclax plus azacitidine, or DMSO control for 15 min at room temperature. After drug incubation, each lysate was divided into twelve 25 µL aliquots and subjected to the same thermal gradient used for intact-cell samples, consisting of 3 min heating at 48–59°C in 1°C increments. Samples were equilibrated at room temperature for 3 min, snap-frozen, and kept frozen until all temperature points were processed. Temperature-point aliquots were pooled by treatment condition and replicate, and soluble proteins were isolated by ultracentrifugation at 100,000 g for 20 min at 4°C. Approximately 220 µL of the soluble supernatant was collected for downstream proteomics.

### Long-term treatments (5 days)

#### Time-course treatment to reach 50% cell viability

For long-term treatment, cells were treated with venetoclax, azacitidine, venetoclax plus azacitidine, or DMSO control under standard culture conditions. Treatments were continued for 5 days, corresponding to the time point at which approximately 50% viability was reached in the VA-treated condition. Cell viability was monitored every 24 h using trypan blue exclusion.

A culture medium change was performed on day 3 to maintain nutrient availability during the long-term treatment. Venetoclax was re-added following medium replacement [45], and azacitidine was replenished every 24 h throughout the experiment to maintain continuous drug exposure. This replenishment strategy was used because azacitidine requires repeated addition during long-term culture [31].

#### Long-term intact-cell CoPISA treatment with Ficoll/Histopaque live-dead separation

After 5 days of treatment, cells from each condition were subjected to Ficoll/Histopaque density-gradient separation to enrich viable cells and separate live and dead fractions. Briefly, cells were resuspended in pre-warmed medium and carefully layered over Ficoll/Histopaque in 15 mL Falcon tubes. Samples were centrifuged at 400 × g for 30 min at room temperature with the centrifuge acceleration and brake turned off to avoid disturbing the density-gradient layers. The live-cell layer was carefully collected, transferred to fresh tubes, washed, and pelleted. Fractions were counted after separation using trypan blue exclusion.

For long-term intact-cell CoPISA profiling, live-cell fractions were processed using the same intact-cell thermal profiling workflow described above. Briefly, cell pellets were washed with PBS, resuspended in 320 µL PBS supplemented with protease inhibitors, and divided into twelve 25 µL aliquots. Aliquots were heated for 3 min across the 48–59°C temperature gradient, equilibrated at room temperature for 3 min, and snap-frozen in liquid nitrogen. Cells were lysed by four freeze–thaw cycles, and temperature-point aliquots were pooled by treatment condition and replicated. Soluble proteins were isolated by ultracentrifugation at 100,000 g for 20 min at 4°C, and the soluble supernatant was collected for proteomics.

#### Long-term expression/abundance module with Ficoll/Histopaque live–dead separation

In parallel with the long-term CoPISA module, SKM-1 cells were treated with venetoclax, azacitidine, venetoclax plus azacitidine, or DMSO control for 5 days and separated into live and dead populations using Ficoll/Histopaque density-gradient separation as described above. No thermal treatment was performed for this module. Instead, live fractions were collected after separation, washed with PBS, and processed directly for expression/abundance-focused proteomics. This module was designed to quantify treatment-associated changes in total protein abundance independently of thermal solubility effects.

### Proteomics Sample Processing

Protein concentrations were measured using a BCA protein assay. For each sample, the volume corresponding to 30 µg of protein was calculated, and sample volumes were normalized using 20 mM 4-(2-hydroxyethyl)-1-piperazinepropanesulfonic acid (EPPS) buffer and transferred to prepared filter tubes (Amicon Ultra-0.5 mL Centrifugal Filters 10KD). Proteins were reduced with 10 mM dithiothreitol for 30 min at 55°C, followed by centrifugation. Reduced cysteines were alkylated with 50 mM iodoacetamide for 60 min at room temperature in the dark. After alkylation, filters were washed with an EPPS buffer to remove excess reagents. Proteins were digested overnight at 37°C using trypsin at an enzyme-to-protein ratio of 1:50. A second digestion was performed the following day using trypsin at 1:100 for 2 h at 37°C. Peptides were recovered by centrifugation, and filters were rinsed with EPPS buffer to maximize peptide recovery.

Before labeling, peptide pH was checked and adjusted to approximately pH 8–8.5. Peptides were labeled using tandem mass tag reagents dissolved in acetonitrile. 16-plex TMT (Thermo Fisher Scientific) reagent was added at approximately fourfold excess relative to peptide amount, with acetonitrile comprising approximately 20–30% of the final labeling reaction. Labeling reactions were vortexed, briefly spun down, and incubated for 2 h at room temperature with shaking. Reactions were quenched by adding hydroxylamine to a final concentration of 0.5%, followed by incubation for 15 min at room temperature. Labeled samples were then combined according to the experimental multiplexing design.

### Peptide fractionation and LC–MS/MS acquisition parameters

TMT-labeled peptides were fractionated by high-pH reversed-phase chromatography using an XBridge Peptide BEH C18 column (3.5 µm, 130 Å, 1 × 150 mm; Waters) on an Ultimate 3000 system (Thermo Scientific). Peptides were separated using an ammonium formate-based gradient at pH 10, and 36 fractions were collected and pooled into 12 fractions using a post-concatenation strategy. The pooled fractions were dried under vacuum and resuspended in 0.1% formic acid prior to LC–MS/MS analysis.

LC–MS/MS analysis was performed using an Orbitrap Eclipse Tribrid mass spectrometer coupled to an Ultimate 3000 nanoLC system and equipped with a FAIMS Pro interface (Thermo Fisher Scientific) and a custom-made column heater set to 60°C. Peptides were separated by reversed-phase chromatography on a C18 analytical column and analyzed in data-dependent acquisition mode. FAIMS compensation voltages of −40 V and −70 V were alternated during acquisition. MS1 spectra were acquired in the Orbitrap, selected precursors were fragmented by CID and MS2 spectra were acquired in the ion trap, and real-time database searching was performed against a human UniProt database. Peptides identified by real-time search were subjected to synchronous precursor selection (SPS)-MS3 analysis, with HCD fragmentation and MS3 acquisition in the Orbitrap for TMT reporter-ion quantification. Fractionation and LC–MS/MS acquisition were performed as previously described [12,46].

### Database Search and Protein Identification

Raw files were processed using MaxQuant version 2.7.0.0, incorporating the Andromeda search engine. Spectra were searched against a reviewed human UniProt protein sequence database (Swiss-Prot) downloaded on 20241010, with a reversed decoy strategy and common contaminants included. Group-specific parameters were used as follows. The type of the search was set to Reporter MS3. Isobaric labels and their correction factors were provided according to the manufacturer’s instructions. Normalization was set to a weighted ratio to the reference channel. Variable modifications were set to oxidation of methionines and acetylation of N-termini, while fixed modification settings contained only carbamidomethylation of cysteines. The maximum number of modifications per peptide was set to 5. Two missed cleavages with trypsin/P were allowed.

Precursor mass tolerances were set to 20 ppm for the first search and 4.5 ppm for the main search, with internal mass recalibration enabled. Peptides of at least 7 amino acids and precursor charge states up to 7 were considered. TMT reporter ion intensities were quantified in Reporter MS3 mode using manufacturer-provided isotope impurity correction factors, and channel intensities were normalized by a weighted ratio to the pooled reference channel. PSM-, peptide-, protein-, and site-level FDRs were controlled at 1% using the target–decoy strategy. Protein identifications require at least one razor or unique peptide.

### Statistical Analysis

Proteomics data were analyzed from primary peptide-to-spectrum match (PSM) output files generated by MaxQuant. Reverse database hits and potential contaminants were removed prior to downstream analysis, and PSMs lacking reporter ion signal across all TMT channels were excluded. Corrected TMT reporter ion intensities were extracted and matched to experimental metadata using sample annotation tables. For each TMT experiment, assay matrices were constructed within the QFeatures framework [47] with PSMs represented as rows and TMT channels represented as columns annotated by treatment condition, biological replicate, and experimental batch.

Reporter ion intensities were normalized using a multistep workflow. Briefly, intensities were converted to relative abundance values by dividing each channel by its total reporter ion signal and scaling to one million to correct for differences in sample loading and total ion intensity. Data were subsequently log2-transformed and quantile normalized to minimize technical variation between samples. Feature aggregation was performed sequentially from PSM to peptide and from peptide to protein level using median summarization. Peptide abundances were obtained by aggregating PSMs sharing identical peptide sequences, followed by protein-level aggregation based on shared protein identifiers. Median-based summarization was implemented using functions from the matrixStats package [48].

Differential protein solubility analysis was performed using the DEP2 framework [49]. Protein-level normalized datasets were represented as SummarizedExperiment objects [50] containing treatment annotations and biological replicate information within the sample metadata. Missing values were imputed using the MinProb method, and pooled reference samples were excluded prior to statistical testing. Differential analysis was carried out using a control-based design in which each treatment condition (A, V, and VA) was compared against the control condition (C). Statistical significance was determined using moderated statistics implemented within the DEP workflow, and proteins were considered significant at a nominal P-value threshold of 0.05. For the long-term living-cell CoPISA dataset, treatment-induced solubility changes were further integrated with protein abundance changes measured in the corresponding long-term expression proteomics dataset. Specifically, log₂ fold changes for each treatment comparison (A/C, V/C, and VA/C) were adjusted using the corresponding abundance-derived log₂ fold changes to generate a derived abundance-corrected solubility score. This metric is intended as a composite index that partially accounts for expression-driven shifts in apparent solubility, while recognizing that both solubility and abundance estimates contain measurement uncertainty and that the resulting transformed values inherit error from both sources. Accordingly, these abundance-adjusted values were interpreted with caution and used primarily for relative prioritization of candidates rather than as absolute measures of protein stability. Both statistical significance and effect size were evaluated during candidate prioritization, with fold-change thresholds used primarily to aid biological interpretation rather than as strict selection criteria.

Given the complexity of comparing single-agent and combination treatments within the CoPISA workflow [46], together with the absence of an established *a priori* error model for low-powered proteomics datasets, we evaluated the robustness of candidate selection using a permutation-based empirical error control strategy. Protein abundance values were randomly permuted across sample labels within each dataset to generate 10,000 randomized null datasets. The identical statistical testing pipeline and selection criteria used for the original data were then applied to each permuted dataset. This approach generated an empirical null distribution of significant hits, enabling assessment of the reproducibility of proteins exhibiting significant alterations [12].

To integrate findings across experimental platforms, a comparative meta-analysis was performed using four independent proteomics datasets: long-term living cell CoPISA, short-term living cell CoPISA, short-term cell lysate CoPISA, and long-term expression proteomics. Only proteins quantified across all datasets were retained for integrative analysis. Differential analysis outputs were merged using common gene identifiers, and treatment-specific P-values, together with log2 fold changes, were extracted for comparative evaluation. Combined statistical significance across datasets was assessed using Fisher’s method for combining independent P-values [25]. Pairwise Fisher combined P-values were calculated for each treatment comparison to integrate evidence across proteomics modalities. Concordance of effect size between datasets was quantified using the geometric mean of absolute log2 fold changes. Proteins were classified as significant using a combined threshold of Fisher p-value < 0.05 together with an effect-size cutoff defined as the upper quartile of the geometric mean absolute fold-change distribution.

## Data Availability

The mass spectrometry proteomics data generated in this study have been deposited in the ProteomeXchange Consortium [51] via the PRIDE partner repository [52] with the dataset identifier PXD084405 and 10.6019/PXD084405 [http://proteomecentral.proteomexchange.org/cgi/GetDataset?ID=PXD084405]. Source data are provided with this paper. There are no restrictions on data availability.

## Code Availability

The analysis code used in this study is publicly available at https://github.com/jafarilab/CoPISA)_long.

## Acknowledgements

This study was financially supported by the Tampere Institute for Advanced Study and the Jane and Aatos Erkko Foundation [Grant 220031 to M.J.]. A.A.S. acknowledges funding from the Swedish Cancer Society (24 3595 Pj), the Swedish Research Council (2023-02692), Åke Wibergs Stiftelse (M23-0186) and Jeanssons Stiftelse (J2023-0094).

## Author Contributions Statement

M.J. conceived the study and designed the research questions. M.J. and A.A.S. provided scientific direction, supervised the project, and contributed to the manuscript preparation. E.G. led the experimental analyses and data interpretation, while M.J. led the computational analyses and data visualization. U.V. participated in proteomics data analysis, M.V. led the validation experiments, and D.R. conducted the proteomics mass spectrometry experiments and data analysis. H.K., K.K.J., M.K., M.V., R.I., B.G., and R.R. supported the interpretation of findings and the validation of the experimental design. E.K. and P.A.H. offered valuable inputs on data interpretation and visualization.

## Supplementary Figures

**Supplementary Fig. 1:**
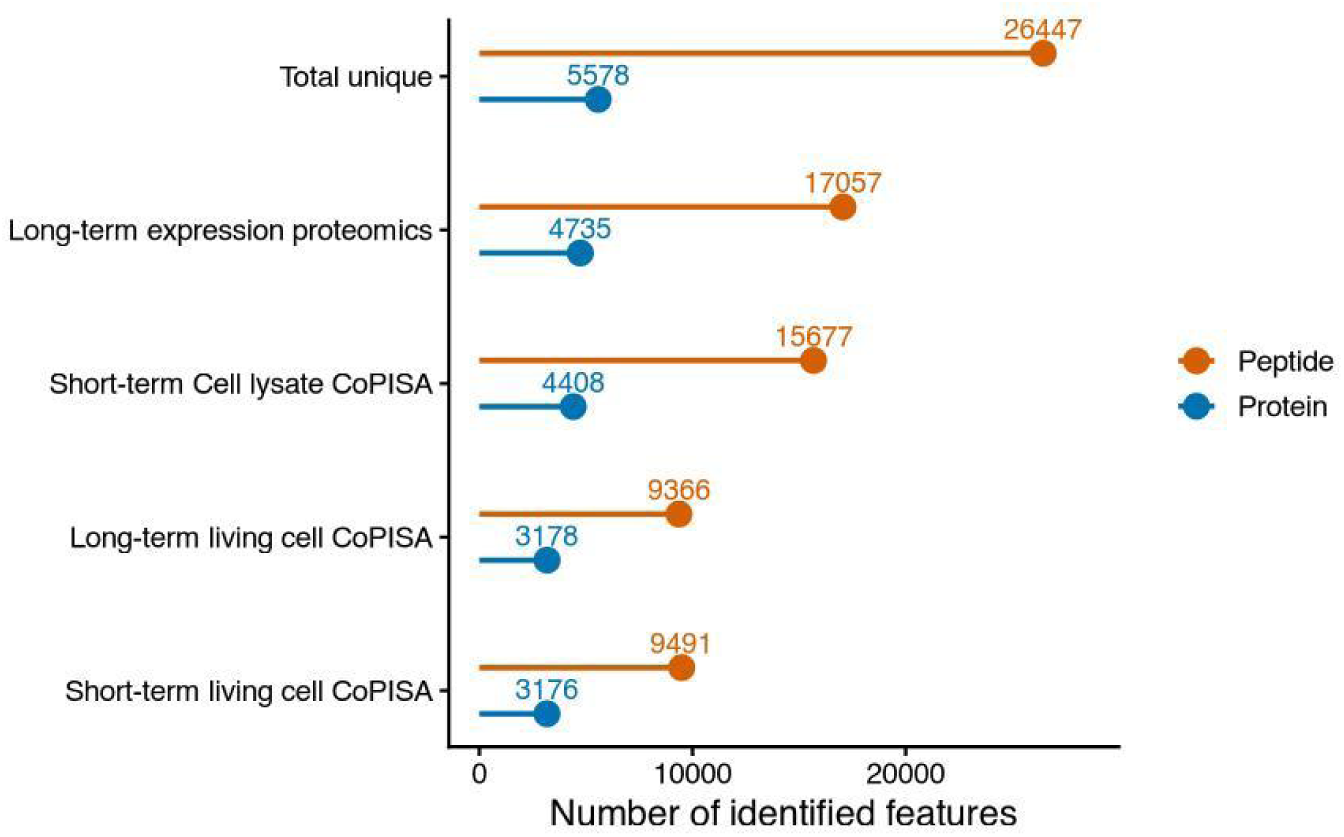
Summary of identified protein and peptide features across all experimental conditions and proteomics workflows. The number of detected features is shown for protein (blue) and peptide (orange) identifications across short-term and long-term living cell CoPISA experiments, short-term cell lysate CoPISA, and long-term expression proteomics. Each condition is additionally compared to the total number of unique proteins and peptides identified across all assays. Feature counts represent the number of non-missing observations per assay, with the final category summarizing the union of all identifications across datasets.

**Supplementary Fig 2:**
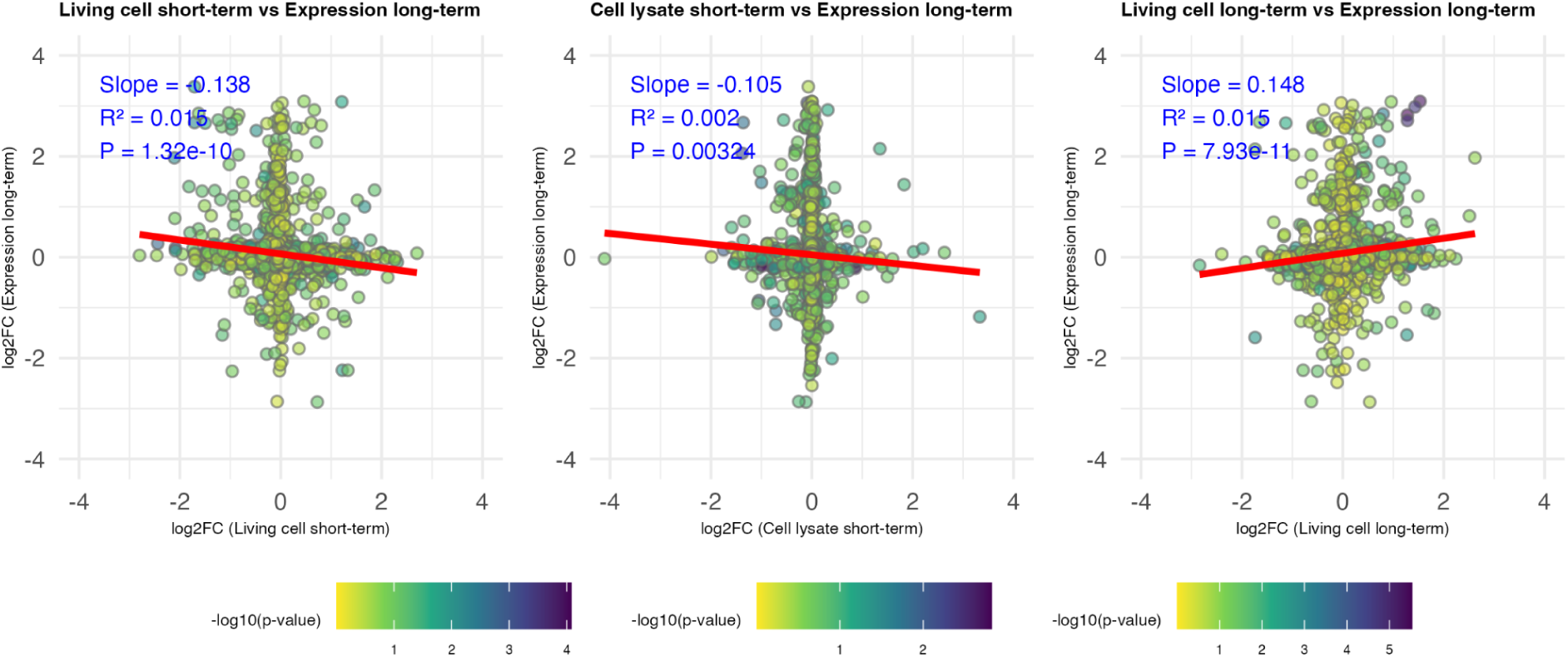
Pairwise fold-change comparisons between CoPISA assays and long-term expression proteomics for the VA combination treatment. FC–FC plots show shared protein identifiers with corresponding log₂ fold-change values for the VA combination treatment relative to control across CoPISA assays and the long-term expression proteomics dataset. Each panel compares one CoPISA-derived fold-change metric against the long-term expression proteomics fold change, together with the p-value from the statistical differential analysis of the long-term expression proteomics dataset. For each comparison, the linear regression coefficient, associated p-value, and fitted regression slope are reported. Source data are provided in the accompanying Source Data file.

**Supplementary File S1. Gene Ontology (Biological Process) enrichment analysis of the retained CoPISA target classes R1 and R3.** Enrichment was performed using Enrichr on the gene sets corresponding to the R1 (CoPISA signal retained, not increased in long-term abundance; n = 35 genes across enriched terms) and R3 (CoPISA signal lost, not increased in long-term abundance; n = 14 genes across enriched terms) classes defined in this study. Each sheet lists, for one class, all tested GO Biological Process terms with the term name and GO identifier, gene overlap (input genes / term gene-set size), nominal P-value, Benjamini–Hochberg-adjusted P-value, Enrichr odds ratio, combined score, and the overlapping input genes.

